# GTL1 is required for a robust root hair growth response to avoid nutrient overloading

**DOI:** 10.1101/2021.06.27.450011

**Authors:** Michitaro Shibata, David S. Favero, Ryu Takebayashi, Ayako Kawamura, Bart Rymen, Yoichiroh Hosokawa, Keiko Sugimoto

**Author notes:** **Author for correspondence:** Michitaro Shibata, Keiko Sugimoto.

## Abstract

- Root hair growth is tuned in response to the environment surrounding plants. While most of previous studies focused on the enhancement of root hair growth during nutrient starvation, few studies investigated the root hair response in the presence of excess nutrients.
- We report that the post-embryonic growth of wild-type Arabidopsis plants is strongly suppressed with increasing nutrient availability, particularly in the case of root hair growth. We further used gene expression profiling to analyze how excess nutrient availability affects root hair growth, and found that RHD6 subfamily genes, which are positive regulators of root hair growth, are down-regulated in this condition.
- On the other hand, defects in GTL1 and DF1, which are negative regulators of root hair growth, cause frail and swollen root hairs to form when excess nutrients are supplied. Additionally, we observed that the RHD6 subfamily genes are mis-expressed in *gtl1-1 df1-1*. Furthermore, overexpression of *RSL4*, an *RHD6* subfamily gene, induces swollen root hairs in the face of a nutrient overload, while mutation of *RSL4* in *gtl1-1 df1-1* restore root hair swelling phenotype.
- In conclusion, our data suggest that GTL1 and DF1 prevent unnecessary root hair formation by repressing *RSL4* under excess nutrient conditions.

## Introduction

Plant biomass accumulation is generally limited by the availability of nutrients, particularly that of the three primary nutrients, nitrogen (N), inorganic phosphate (Pi) and potassium (K). Therefore, fertilizer application is an effective means of increasing plant growth in conditions where nutrients are otherwise lacking (Barker & Pilbeam, 2015). Fertilizers supported the past green revolution and have played an essential role increasing the world food supply (Herdt & Capule, 1983). However, because crops are generally not seriously affected by excess nutrients but produce lower yields in the absence of sufficient nutrients, fertilizers are often applied in excess (Barker & Pilbeam, 2015). Arabidopsis, the most widely used model plant, also does not typically exhibit obvious growth differences in the presence of excess nutrients, except in the cases of certain heavy metals, B and N sources (Li *et al*., 2005; Gruber *et al*., 2013; Esteban *et al*., 2016; Yoshinari & Takano, 2017). Thus, plant responses to excess nutrients have been largely overlooked, and the mechanisms plants use to protect themselves from excess nutrition are largely unknown.

Root hairs, which grow from the root epidermis, impact nutrient uptake from the soil, as they increase the surface area of the root system. In Arabidopsis, root hairs generally initiate from trichoblasts, one of two types of epidermal cells specified during root development. Atrichoblasts, on the other hand, are root epidermal cells that do not produce hairs under normal conditions (Ishida *et al*., 2008; Grierson *et al*., 2014; Salazar-Henao *et al*., 2016). Root hair initiation, as well as growth of root hairs, is precisely regulated based on nutrient availability. For instance, Pi starvation promotes both root hair formation and growth. Under Pi starvation, root hair number is increased by the initiation of root hairs from atrichoblasts and also by the production of multiple hairs from a single epidermal cell (Ma *et al*., 2001). At the same time, root hair length increases nearly three-fold in low compared to high phosphate conditions (Bates & Lynch, 1996). Iron deficiency also strongly affects root hair development (Schmidt *et al*., 2000). In contrast to Pi starvation, however, iron deficiency increases root hair branching rather than promoting ectopic root hair formation (Müller & Schmidt, 2004). These findings suggest that at least some environmental signals affect root hair development independently of each other.

A number of transcription factors have been identified that play key roles regulating root hair development (Ishida *et al*., 2008; Bruex *et al*., 2012; Salazar-Henao *et al*., 2016; Franciosini *et al*., 2017; Shibata & Sugimoto, 2019; Vissenberg *et al*., 2020). Atrichoblasts are characterized specifically by the expression of *GLABRA2* (*GL2*), which encodes a transcription factor that functions as a negative regulator of root hair formation and is often used as a marker for non-hair cells (Masucci *et al*., 1996; Di Cristina *et al*., 1996; Lin *et al*., 2015). Conversely, the bHLH transcription factors ROOT HAIR DEFECTIVE6 (RHD6) and RHD6-LIKE1 (RSL1) are key factors that promote hair development (Masucci & Schiefelbein, 1994; Menand *et al*., 2007; Pires *et al*., 2013). Following specification of root epidermal cells as trichoblasts, RHD6 and RSL1 promote transcription of genes encoding other bHLH transcription factors, including *RSL2*, *RSL4* and *Lotus japonicus ROOTHAIRLESS-LIKE 3* (*LRL3*) (Masucci & Schiefelbein, 1994; Karas *et al*., 2009; Yi *et al*., 2010). The *rsl2 rsl4* double mutant completely lacks root hairs (Yi *et al*., 2010), indicating that *RSL2* and/or *RSL4* are essential for root hair growth. Similarly, the double or triple mutants for *LRL3* and its homologues *LRL1* and/or *LRL2* have short root hairs that occur at a lower density than normal (Karas *et al*., 2009; Tam *et al*., 2015; Breuninger *et al*., 2016). Thus LRL3, together with LRL1 and LRL2, contributes to both root hair formation and growth. In addition to promoting growth of root hairs in response to developmental signals, bHLH transcription factors, particularly RSL2 and RSL4, are also important for root hair growth induced by exogenous phytohormone treatments (auxin, cytokinin, ethylene or jasmonic acid) or nutrient (Pi, N or Fe) deficiency (Yi *et al*., 2010; Datta *et al*., 2015; Zhang *et al*., 2016; Mangano *et al*., 2017; Feng *et al*., 2017; Bhosale *et al*., 2018; Han *et al*., 2020b). Therefore, RSL2 and RSL4 appear to be core factors in the gene regulatory network (GRN) that controls root hair growth (Lee & Cho, 2013; Franciosini *et al*., 2017; Shibata & Sugimoto, 2019). In addition to these positive regulators of root hair growth, negative regulators have also been identified. The trihelix transcription factor GT2-LIKE1 (GTL1) and its closest homolog, DF1, terminate root hair growth by directly repressing *RSL4* together with RSL4 target-genes (Shibata *et al*., 2018). In addition, a DOF-type transcription factor, OBF BINDING PROTEIN 4 (OBP4), is a negative regulator of root hair growth, as induction of *OBP4* reduces root hair length (Rymen *et al*., 2017). Unlike GTL1 and DF1, OBP4 represses *RSL2* expression and does not affect *RSL4* expression, suggesting that plants have at least two transcriptional pathways that repress root hair growth.

In this study we investigated how root hairs are affected by excess nutrients in Arabidopsis. Specifically, we demonstrate that root hair growth is strongly suppressed on double-strength Murashige-Skoog (2xMS) media. Further, we show that the *gtl1-1 df1-1* mutant forms frail root hairs on 2xMS, suggesting that GTL1 and DF1 prevent aberrant root hair development in the presence of excess nutrients. These findings shed light on the mechanisms that plants have evolved to adapt to growth in variable conditions.

## Materials and Methods

### Plant materials and growth conditions

The *gtl1-1* (WiscDsLox413-416C9), *df1-1* (SALK_106258), *rsl4-1*(Yi et al., 2010), *rhd6-3*(GABI-Kat 475E09), *obp4-2* (SALK_08510), *obp4-3* (SALKseq_108296), *pGTL1:GTL1-GFP*, *pEXP7:GTL1-GFP, pGL2:GL2-GFP/gl2-8,* and *pEXP7:NLS-GFP* lines were previously described (Breuer *et al*., 2009, 2012; Yi *et al*., 2010; Ikeuchi *et al*., 2015; Rymen *et al*., 2017; Shibata *et al*., 2018). *lrl3-2* corresponds to SALK_012380 (Fig. S7). *35S:XVE>>RHD6* (CS2104357, (Coego *et al*., 2014) were obtained from ABRC. 10 µM 17β-estradiol were applied to induce the *RHD6*. Both *RSL4* cDNA and genomic DNA overexpression lines were generated for this research.

Plants for most experiments were grown in a MLR-352H-PJ plant growth chamber (PHCbi) with a constant temperature of 22°C, at 60% relative humidity and under continuous light (50-70 µmol/m2/sec). Half, single or double strength MS media with 1% sucrose, 0.55 mM myo-inositol, 2.5mM MES-KOH (pH5.7), and 6 g/l Gelzan (Sigma) were used for plant growth. Each MS medium was made by adding 1 bag of MS salt mix (Nihon pharmaceutical CO., LTD.) into the proper volume of milli-Q (Merck) water. For self-made MS media, each chemical solution was prepared and mixed before autoclaving. FeSO_4_ was added together with 2Na-EDTA to avoid precipitation.

### Plant growth analysis

To measure root hair length, root images were taken from main roots of 12 days old seedlings with a dissection microscope (Leica M165 FC equipped with a digital Leica DFC 7000T camera). For the measurements of main root length and lateral root length, root images from 14 days old seedlings were taken using a scanner (Epson GT-X830). Subsequently, the shoot tissues from the same seedlings were collected for the measurements of fresh weight. Five seedlings were pooled and weighed on a microbalance (METTLER TOLEDO AT200). All image analysis was manually done with ImageJ (1.53g).

### Microscopic observation

Expression patterns of *pEXP7:NLS-GFP* and *pGL2:GL2-GFP* were examined using a Leica SP5 confocal laser scanning microscope with the Leica HyD hybrid detector. Argon-Ion laser was used for the excitation and the emission light was detected using a spectral detector. The root images were produced from multiple z-stack images with maximum projection on LAS-X software (Leica).

### Plasmid construction and plant transformation

For *35S:RSL4gDNA-GFP* and *35S:RSL4cDNA-GFP*, the coding regions of *RSL4* (AT1G27740.1) were amplified by PCR (PrimeStar max, Takara) using genomic DNA and cDNA as templates, respectively. The PCR products were introduced into pDONR221(Invitrogen) via BP reactions and subsequently transferred to the *pGWB505* destination vector (Nakagawa *et al*., 2007) via LR reactions. The resulting constructs were then introduced into Arabidopsis plants via the floral dip method (Clough & Bent, 1998). A set of primers used for PCR amplification is provided in Table S3.

### Atomic force microscopy (AFM)

Root sections from 1-week-old seedlings were used for measuring root hair strength. The samples were immobilized by molten 1.5% low-melting point agarose (Agarose LMT 1-20K, PrimeGel, Takara Bio Inc.) spread thinly on a glass-bottom dish and immediately covered in water. Intact root hairs were selected for measurement with an inverted microscope (IX71, Olympus). To reduce the effect of leverage, a distance of 30 µm from the root hair base was used as the contact point. Data were obtained using an AFM system (JPK instruments, Nanowizard 4), to which an AFM cantilever (NANOSENSORS, SD-Sphere-NCH, force constant: 42 N/m) was attached. The cantilever tip is shaped into a hemisphere (400 nm radius) to avoid damaging the contact point on the cell. Force curves were observed using contact mode with set point at 130 nN and approach velocity at 2.0 µm/s. The data were analyzed by the Hertz model, using JPK Data Processing (ver 6.1.158) software, in which the apparent elastic modulus of the cell (stiffness score) was estimated by Young’s modulus.

### RNA extraction and RT-qPCR analysis

Total RNA was extracted from 7-day-old roots using the RNeasy Plant Mini Kit (Qiagen) and QIAcube (Qiagen). Extracted RNA was reverse transcribed using a PrimeScript RT-PCR kit with DNase I (Perfect Real Time) (Takara) in accordance with the accompanying protocol. Transcript levels were determined via qPCR using the THUNDERBIRD SYBR qPCR Mix kit (Toyobo) and Mx399P QPCR system (Agilent). The expression of either the *UBQ10* or *HEL* gene was used as an internal control. A set of primers used for RT-qPCR is provided in Table S3.

### Promoter-luciferase assay

The effector vectors *35S:GTL1* and *35S:RHD6* were previously described in Shibata et al., 2018 and Rymen et al. 2017, respectively. For the reporter vectors, the promoter regions were amplified by PCR (PrimeSTAR max, Takara) and cloned into pGEM/T-EASY containing LUC+ for *RSL4* and *LRL3*, and R4L1pUGW35 (Nakamura *et al*., 2009) for *RPS5A* and *RHD6*. Primers used for the cloning are shown in Table S3. As an internal control, the *pPTRL* vector, which drives the expression of a *Renilla LUC* gene under the control of the *CaMV p35S* promoter, was used. The constructs were introduced into Arabidopsis MM2d culture cells (Menges & Murray, 2002) by a gold particle bombardment system. Luciferase activity was quantified using a Mithras LB940 microplate luminometer (Berthold Technologies) as described previously (Hiratsu *et al*., 2002).

## Results

### Excess nutrients inhibit Arabidopsis growth

Murashige-Skoog (MS) medium (Murashige & Skoog, 1962) is widely used for plant growth. For Arabidopsis, half-strength (1/2x) and full-strength (1x) MS media are commonly used as “normal” growth conditions. Thus, in addition to 1/2x and 1x MS media, we prepared double-strength (2x) MS medium as an “excess nutrient” condition to study the impact of excess nutrients on post-embryonic development of Arabidopsis.

As shown in Fig.1, plant growth was severely affected by MS strength. Consistent with the visible effect on growth, the fresh weight of 2-weeks-old seedlings (shoot) significantly decreased with increasing MS strength (Fig. 1b). Although the main root length was not significantly altered between the 1/2x and 1xMS conditions, the total lateral root length obviously declined as MS strength increased (Fig. 1d). This observation is consistent with previous reports describing that plants generally reduce lateral root development in the presence of excess nutrients in order to decrease nutrient uptake (Gruber *et al*., 2013).

**Fig. 1.**
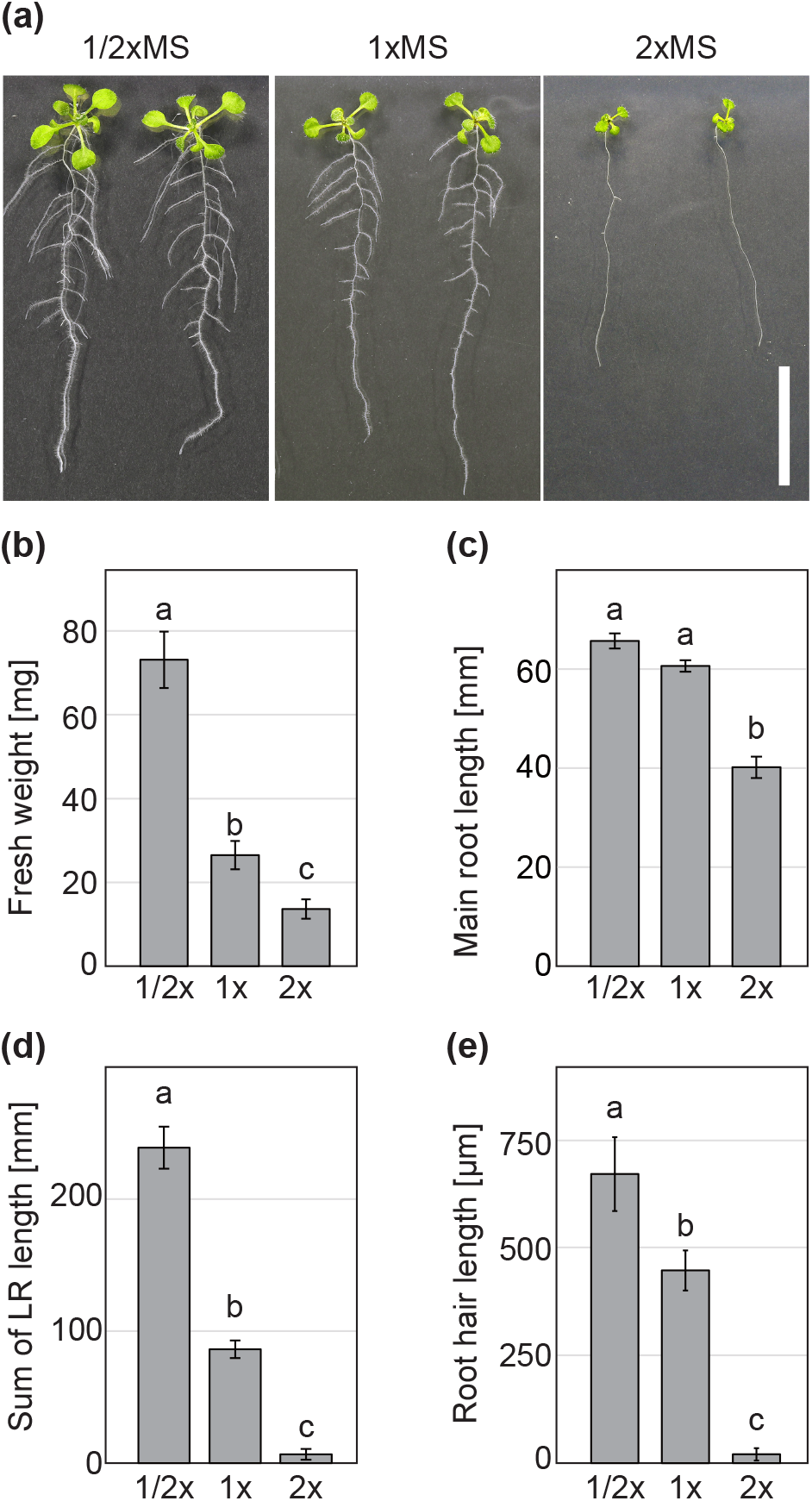
Arabidopsis growth is repressed by higher concentrations of MS. (a) Images of 3-week-old WT plants grown on half- (1/2x), full- (1x) or double- (2x) strength MS medium. Scale bar = 1 cm. Quantitative analysis of fresh weight (b), main root length (c), sum of lateral root (LR) length (d) and length of 20 longest root hairs from each seedling (e). Data are mean± SE (n = 40 for (b), 16 for (c) and d, 12 for (e)). Different letters indicate significant differences One-way ANOVA with post-hoc Tukey HSD test, p < 0.01).

### Double-strength MS medium suppresses root hair growth

In addition to affecting lateral roots, MS concentration also affected root hairs. To study the impact of excess nutrition on root hair development in greater detail, we observed root hairs under high magnification (Fig. 2a, left). Consistent with the observation, the root hair length gradually decreased with increasing MS strength (Fig. 1e). Two weeks after germination, only extremely short root hairs were observed on 2xMS medium. Close inspection of root hairs on 2xMS medium revealed that dome-like structures were established on epidermal cells, indicating that root hair initiation had progressed (Fig. 2b). To confirm the root hair/non-hair cell identities, we observed the expression patterns of *EXPANSIN A7* (*EXPA7*) and *GL2*, which are hair cell and non-hair cell markers, respectively (Masucci *et al*., 1996; Cho & Cosgrove, 2002). In the case of Arabidopsis, since hair cells and non-hair cells are tandemly arranged within cell files in the root epidermis, GFP signals from both *pEXP7:NLS-GFP* and *pGL2:GL2-GFP* were observed in strips (Fig. 2c,d). On 2xMS medium, each GFP signal also aligned within cell files, indicating that root hair and non-hair cell-pattern was conserved (Fig. 2c,d). In addition, we also examined the expression levels of *EXPA7* and *GL2* (Fig. S1). For *EXPA7*, the expression level decreased on 2xMS. This is reasonable because *EXPA7* contributes to cell expansion (Cho & Cosgrove, 2002), and does not necessarily imply that there are fewer hair cells in this condition. Importantly, *GL2* expression was also reduced on 2xMS (Fig. S1). This suggests that the extreme short root hair response is not caused by an increase of non-hair cells in the root epidermis due to an increase of the GL2 root hair identity repressor. Taken together, these results indicate that the root hair phenotype on 2xMS media is caused by defects in cell growth rather than cell specification.

**Fig. 2.**
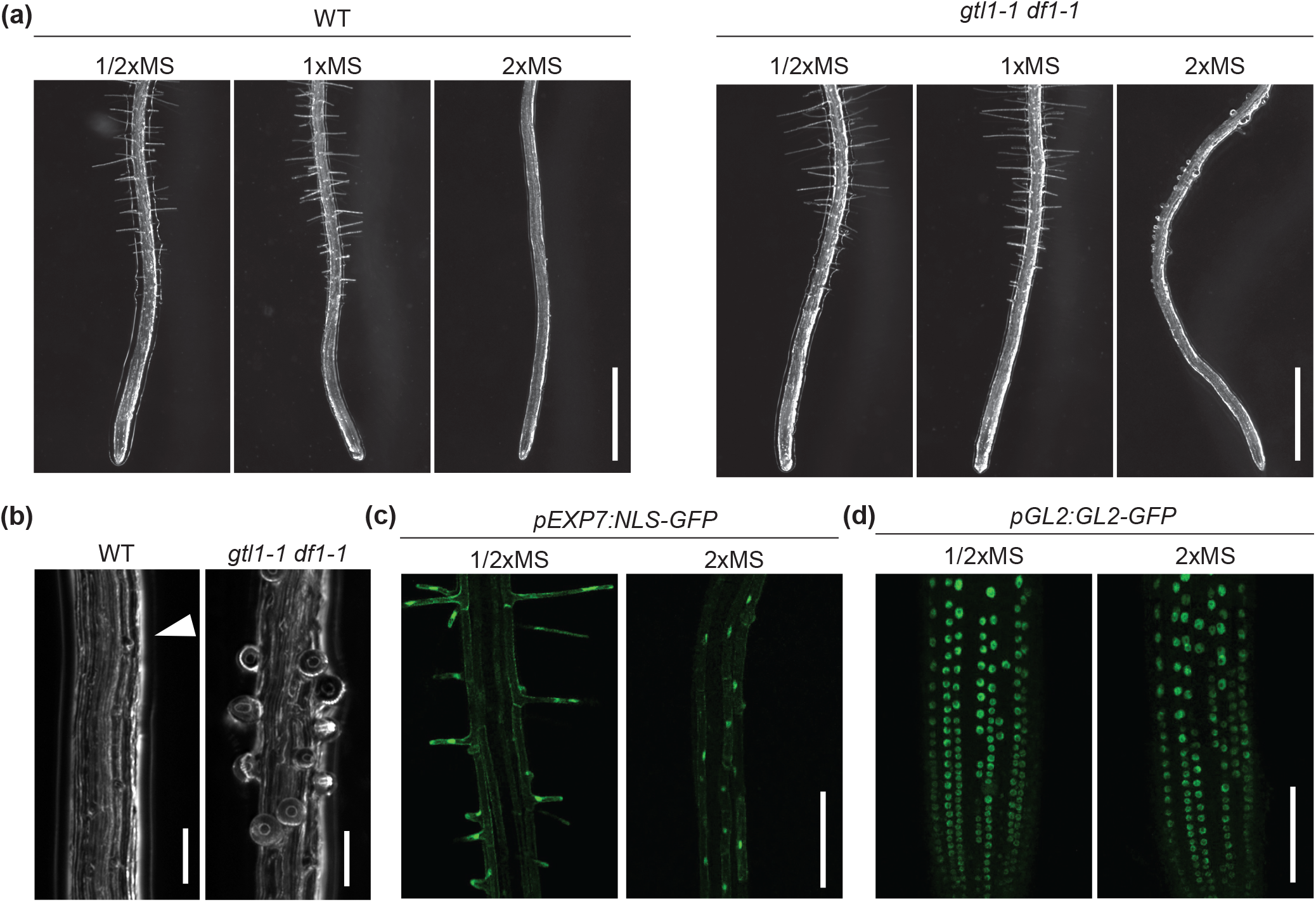
GTL1/DF1 prevent formation of aberrant root hairs on 2xMS by repressing root hair growth. (a) Images of root tips of WT and *gtl1-1 df1-1* on half- (1/2x), full- (1x) or double- (2x) strength MS medium. High-magnification images of roots grown on 2xMS media are shown in (b). The arrow head in (b) shows a root hair initiation site from an epidermal cell. Scale bars = 1000 µm in (a) and 100 µm in (b). (c, d) Confocal microscope images of *pEXP7:NLS-GFP* (c) and *pGL2:GL2GFP/gl2-8* (*pGL2:GL2-GFP*) (d) on 1/2xMS and 2xMS. *EXP7* and *GL2* are used as hair cell and non-hair cell markers, respectively. Scale bars = 200 µm in (d) and 100 µm in (e).

We previously identified the trihelix transcription factors *GTL1* and *DF1* as negative regulators of root hair growth that do not affect root hair cell fate (Shibata et al., 2018). Therefore, we hypothesized that GTL1 and DF1 might contribute to the root hair phenotype on 2xMS medium. To test this hypothesis, we grew the *gtl1-1 df1-1* mutant on 1/2x, 1x, and 2xMS media and observed the root hair phenotypes (Fig. 2b, right). On 1/2x- and 1xMS media, the *gtl1-1 df1-1* mutant had longer root hairs compared to those of the WT, which is consistent with our previous data (Shibata et al., 2018). Strikingly, in the presence of 2xMS, we found that *gtl1-1 df1-1* formed tiny root hairs that were more obvious than the dome-like structures observed in the WT (Fig. 2b). Also, in contrast to the WT, most root hairs found in *gtl1-1 df1-1* grown on 2xMS were swollen and a few hairs had ruptured (Fig. 2b). These results suggest that GTL1 and DF1 are required for proper termination of root hair growth and/or maintenance of the tube-like structure of root hairs on 2xMS medium.

### Swollen root hairs lose stiffness

In order to further investigate the morphological differences between WT and *gtl1-1 df1-1* root hairs, we estimated the stiffness scores of root hairs from analysis of the force-indentation curves acquired by the AFM measurement. First, we confirmed root hair stiffness in plants grown in a normal nutrient condition, i.e. on 1/2xMS medium. In the case of the 1/2xMS condition, no significant difference in the AFM measurement was detected between the WT and *gtl1-1 df1-1* (Fig. 3a,c, S2). The resulting score for the WT was 4.04 ± 3.31 MPa, which was higher than the score obtained from main roots (0.47 ± 0.12 MPa) (Fig. S2b).

**Fig. 3.**
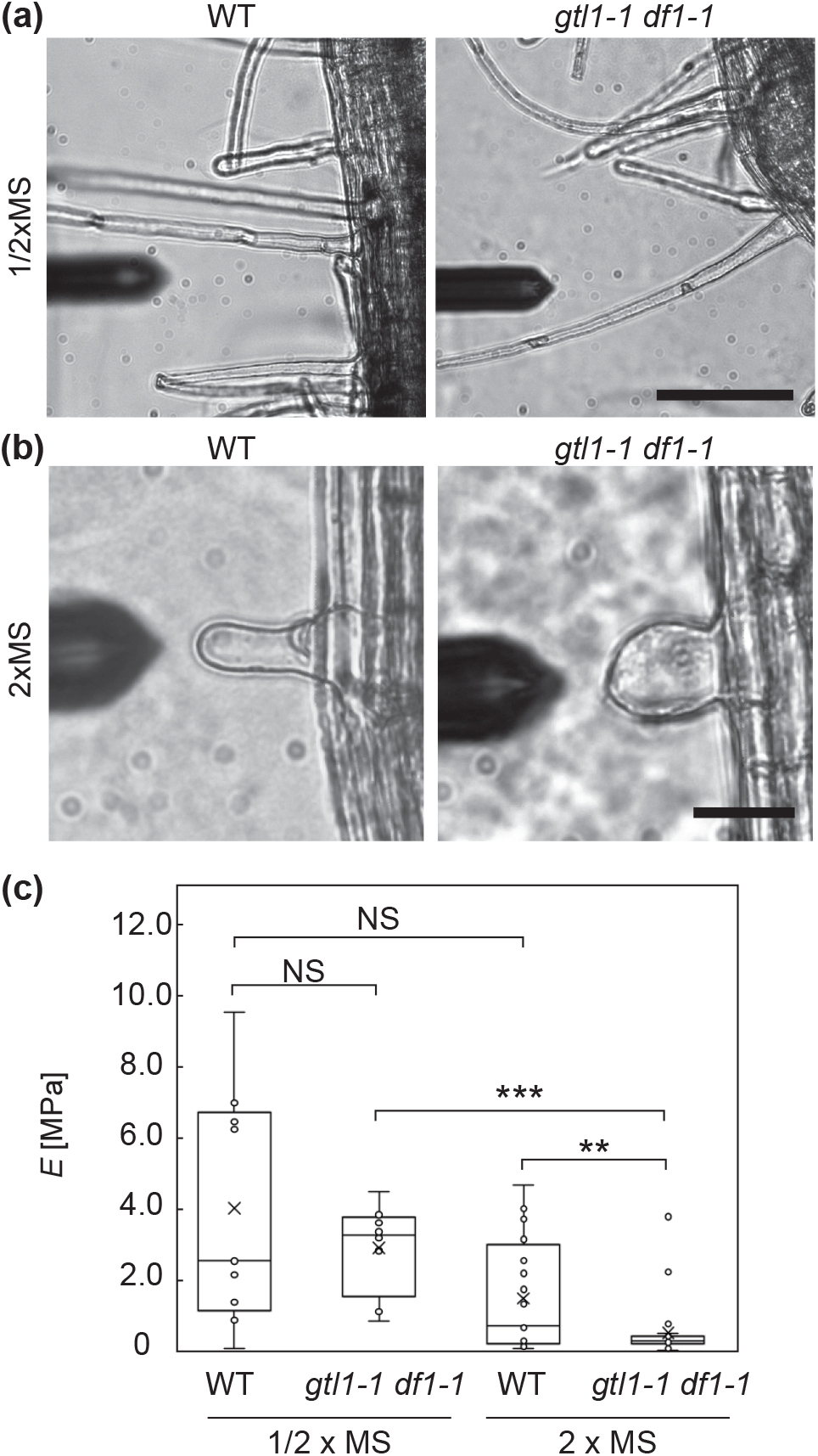
GTL1/DF1 maintain root hair stiffness on 2xMS. (a,b) Microscopic images of root hairs used for AFM analysis. The cantilever indicates the root hairs used for measurement. Scale bars = 100 µm in (a) and 30 µm in (b). (c) Box plots showing root hair stiffness scores, as determined from AFM analysis. Y axis indicates Young’s modulus (*E*). Asterisks indicate a significant difference (Student’s t-test, *p < 0.05, **p < 0.01 ***p < 0.001).

On the contrary, for the 2xMS condition, an obvious difference in root hair stiffness was observed in *gtl1-1 df1-1* compared to the WT (Fig. 3b,d). More specifically, the majority of root hairs in WT were stiff, while a few were loose. In contrast, most *gtl1-1 df1-1* root hairs were loose and only a few were stiff (Fig. 3d). Compared to main roots, root hairs in the WT were much stiffer, whereas root hairs and main roots produced similar stiffness readings in *gtl1-1 df1-1* (Fig. S2b). These data indicate that, besides being swollen, the root hairs in *gtl1-1 df1-1* have lost a mechanical property typically associated with this tissue.

### A combination of multiple nutrients alters root hair growth

To identify which nutrient(s) cause the strong suppression of root hair growth on 2xMS medium, we prepared custom MS media and altered the ingredients. First, we prepared a medium named “2xMS_All”, which mimics 2xMS made from an MS salt mix (Table S1), and confirmed that root hair growth was strongly inhibited in WT plants grown on this medium (Fig. 4a). Next, to identify which nutrients cause the strong suppression of root hair growth, we prepared various MS media, each in which one ingredient is reduced from the 2xMS level to that of 1/2xMS (Table S2). Among 14 different chemicals found in MS salts, we found that the reduction of NH_4_NO_3_ alone (2xMS_1/2xNH_4_NO_3_) partially restored root hair growth (Fig. 4a). Others did not visibly affect root hair growth on 2xMS. Further, to evaluate the impact of N source on root hair growth, we prepared medium with reduced KNO_3_ and NH_4_NO_3_. Although reduction of only KNO_3_ (2xMS_1/2xKNO_3_) did not restore root hair growth, decreased levels of both KNO_3_ and NH_4_NO_3_ (2xMS_1/2xKNO_3_ 1/2xNH_4_NO_3_) further restored root hair growth compared to 2xMS_1/2xNH_4_NO_3_ (Fig. 4a).

**Fig. 4.**
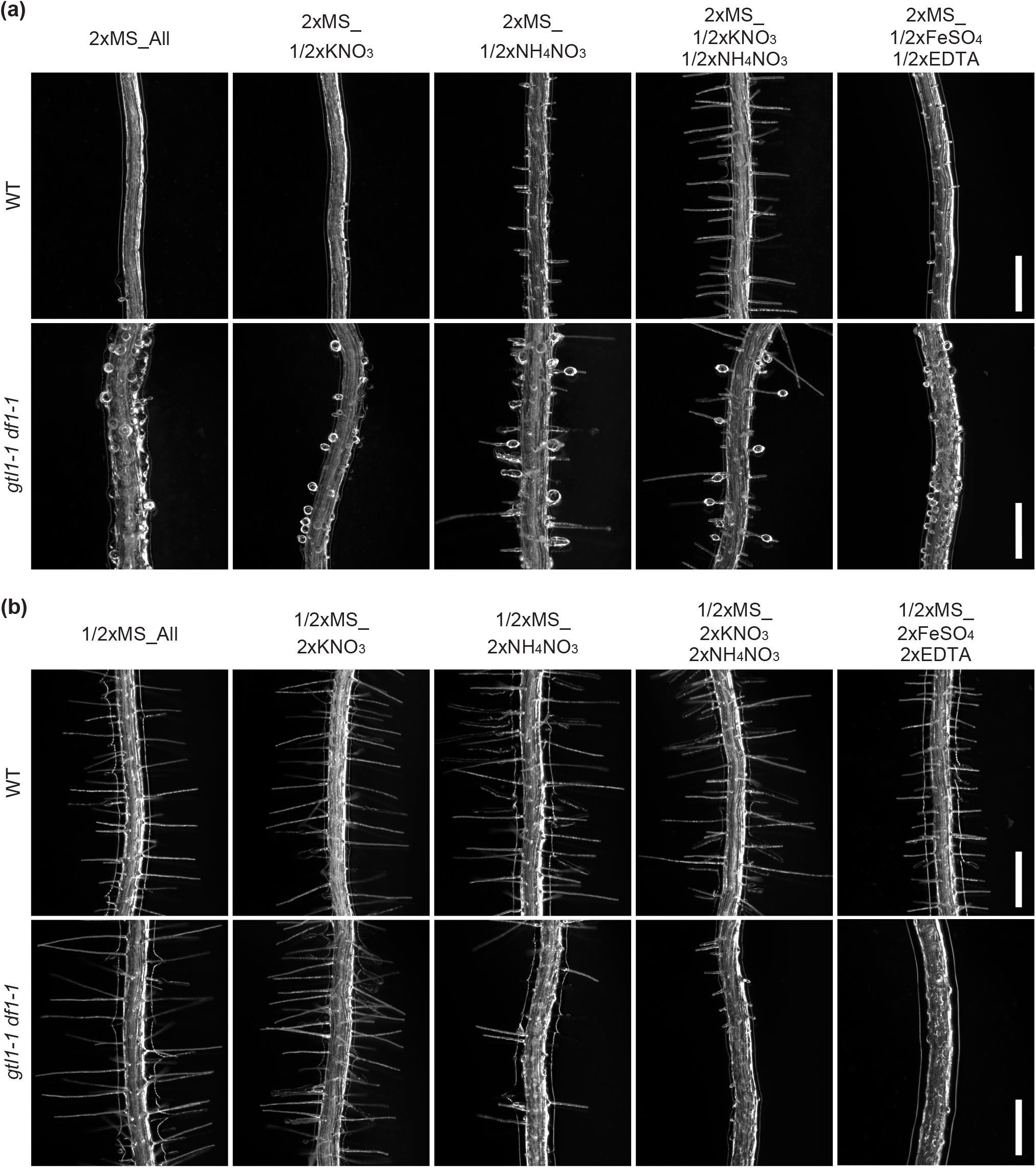
Multiple nutrients contained in MS media affect root hair growth. Images of root hairs grown on custom MS media. 2xMS- and 1/2xMS-based medium are shown in (a) and (b), respectively. Upper and lower panels show root hairs of WT and *gtl1-1 df1-1* in each condition, respectively. See Table S2 for a complete list of ingredients. Scale bars = 200 µm.

Additionally, to investigate if double-strength KNO_3_ and/or NH_4_NO_3_ are sufficient to suppress root hair formation, we prepared custom 1/2xMS with extra nutrients (Table S2). First, we confirmed that WT plants form normal root hairs on our self-made 1/2xMS medium (1/2xMS_All, Fig. 4b). Next, we observed root hair phenotypes in plants grown on 1/2xMS media with 2xMS levels of KNO_3_ (1/2xMS_2xKNO_3_), NH_4_NO_3_ (1/2xMS_2xNH_4_NO_3_) or both KNO_3_ and NH_4_NO_3_ (1/2xMS_2xKNO_3_ 2xNH_4_NO_3_). As opposed to reduction of these nutrients in 2xMS, addition of these nutrients to 1/2xMS did not clearly affect root hair growth. Instead, we found that double-strength iron (1/2xMS_2xFeSO4_4_+2xEDTA) strongly reduced root hair growth (Fig. 4b), although reduction of the iron source in 2xMS (2xMS_1/2xFeSO4_4_+1/2xEDTA) had little effect on root hair growth (Fig 4a). The response of root hairs to double-strength iron is consistent with previous studies demonstrating that root hair growth is sensitive to iron levels (Schmidt *et al*., 2000). It is important to note, however, that double-strength iron or N alone did not reproduce the root hair phenotype observed on 2xMS_All. Therefore, we conclude that the effect of 2xMS on root hair growth is likely caused by a combination of multiple nutrients, including N and Fe.

In a similar manner, we studied which nutrients cause the root hair swelling in g*tl1-1 df1-1* on 2xMS. First, we confirmed that *gtl1-1 df1-1* forms swollen root hairs on our self-made 2xMS medium (Fig. 4a). Next, we observed root hair morphology of *gtl1-1 df1-1* on altered 2xMS media, and found that reduction of NH_4_NO_3_ slightly mitigated the root hair swelling phenotype in *gtl1-1 df1-1* (Fig. 4a). Some of the root hairs of *gtl1-1 df1-1* grown on 2xMS_1/2xNH_4_NO_3_ were tube-like rather than balloon-like in structure. We next observed growth of the mutant on 2xMS medium with reduced KNO_3_ and NH_4_NO_3_ (2xMS_1/2xKNO_3_ 1/2xNH_4_NO_3_). As shown in Fig. 4a, the root hair-swelling phenotype in *gtl1-1 df1-1* was further rescued by the reduction of both N sources. However, the majority of root hairs were still partially swollen, which means that the reduction of N alone does not fully restore the root hair phenotype.

Furthermore, in order to study if double-strength KNO_3_ and/or NH_4_NO_3_ in 1/2x MS are sufficient to induce root hair swelling in *gtl1-1 df1-1*, we again used our custom MS media series (Table S2). First, we confirmed that *gtl1-1 df1-1* forms normal root hairs on our self-made 1/2xMS (1/2xMS_All, Fig 4a), and then observed root hair phenotypes on the 1/2xMS media containing extra nutrients. Consistent with both Johnson and commercial MS media, *gtl1-1 df1-1* had longer root hairs on self-made 1/2xMS compared to the WT. On the contrary, root hair formation of *gtl1-1 df1-1* was strongly inhibited by addition of NH_4_NO_3_ to 1/2xMS (1/2xMS_2xNH_4_NO_3_). Moreover, addition of KNO_3_ to 1/2xMS with double-strength 2x NH_4_NO_3_ (1/2xMS_2xKNO_3_ 2xNH_4_NO_3_) further inhibited root hair formation. In addition, double-strength iron (1/2xMS_2xFeSO4_4_+2xEDTA) completely suppressed root hair growth in *gtl1-1 df1-1* (Fig. 4b). Notably, such a severe phenotype was only observed in *gtl1-1 df1-1* grown on 1/2xMS_2xFeSO4_4_+2xEDTA and not in the WT. These results suggest that *gtl1-1 df1-1* is hypersensitive to environmental effects on root hair growth. However, from these experiments, we were unable to identify the specific ingredients in 2xMS that trigger root hair swelling. This likely indicates that a combination of multiple nutrients causes root hair swelling in *gtl1-1 df1-1*.

### *RHD6*, *RSL4* and *LRL3* are differentially expressed in *gtl1-1 df1-1*

Based on the root hair phenotypes, we concluded that WT plants suppress root hair growth on 2xMS via the activity of GTL1 and DF1. Thus, we hypothesized that *GTL1* and *DF1* are induced by higher MS concentrations. In order to test this hypothesis, we measured mRNA levels of *GTL1* and *DF1* by RT-qPCR. However, contrary to expectations, the expression of both *GTL1* and *DF1* actually declined on 2xMS compared to 1/2xMS (Fig. S3), suggesting that activity of GTL1 and DF1 is regulated by non-transcriptional mechanisms.

We next investigated the molecular mechanisms by which the WT suppresses root hair growth on 2xMS and how defects in *GTL1* and *DF1* cause root hair swelling on 2xMS. Since members of the RHD6 subfamily generally play key roles promoting root hair growth in response to environmental signals (Franciosini *et al*., 2017; Shibata & Sugimoto, 2019), we investigated if the expression of the genes encoding these transcription factors is affected by MS strength. Between the WT and *gtl1-1 df1-1*, we found that *RHD6*, *RSL2*, *RSL3*, *RSL4* and *LRL3* were differentially expressed in at least one condition (Fig. 5). Among these genes, *RHD6*, *RSL4* and *LRL3* were expressed at higher levels in *gtl1-1 df1-1*, which is consistent with our previous data (Shibata *et al*., 2018). Interestingly, the expression of *RHD6* and *RSL4* was reduced on 2xMS compared to both 1/2xMS and 1xMS. Additionally, the expression levels of these genes in *gtl1-1 df1-1* on 2xMS were similar to that in the WT on 1/2xMS or 1xMS, suggesting that the suppression of *RHD6* and *RSL4* expression on 2xMS is deficient in *gtl1-1 df1-1*. In contrast to *RHD6* and *RSL4*, *RSL2* and *RSL3* levels were lower in *gtl1-1 df1-1* compared to the WT in at least one condition. Notably, the expression of both *RSL2* and *RSL3* was significantly reduced in *gtl1-1 df1-1* on 2xMS. These data indicate that *gtl1-1 df1-1* causes stronger suppression of *RSL2* and *RSL3* expression on 2xMS.

**Fig. 5.**
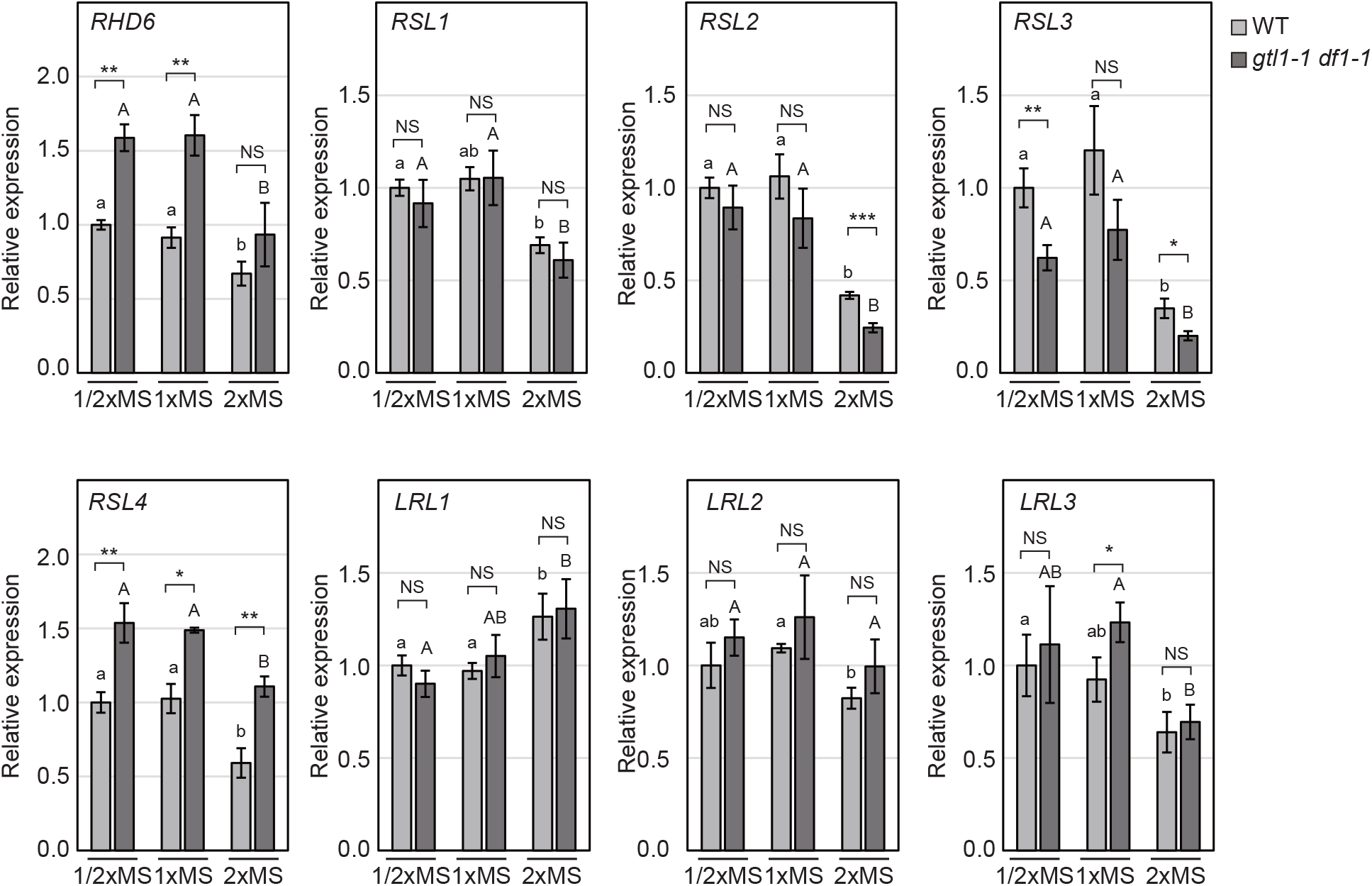
The expression of root hair regulators is affected by MS concentration. RT-qPCR analysis of key transcription factors regulating root hair growth in WT and *gtl1-1 df1-1* grown on 1/2x, 1x and 2x MS media. Expression levels are normalized to that of the *HEL* gene. Data are mean +/- SD. (n = 3, biological replicates). Asterisks indicate a significant difference compared to the WT grown on the same type of medium (Student’s t-test, *p < 0.05, **p < 0.01). Different letters indicate significant differences between media conditions within the same genotype (One-way ANOVA with post-hoc Tukey HSD test, p < 0.05).

In addition to *RHD6* and *RSL4* mentioned above, the levels of *RSL2, RSL3*, and *LRL3* declined on 2xMS compared to other media in the WT (Fig. 5). These expression profiles are consistent with the near lack of root hair growth in the WT on 2xMS. The *gtl1-1 df1-1* mutant also exhibited reduced expression of these *RHD6* subfamily genes on 2xMS, demonstrating that other suppression mechanisms are still active. A Dof-type transcription factor OBP4 was characterized as a suppressor of *RSL2* and *RSL3* that inhibits root hair growth in response ABA signaling (Rymen *et al*., 2017). To study whether OBP4 is required for the repression of *RSL2* and *RSL3* on 2xMS, we analyzed *obp4-2* (SALK_085101, knock-down allele) and *obp4-3* (SALKseq_108296, knock-out allele) mutants across the MS dilution series. The roots of *obp4-2* and *obp4-3* resembled the WT on 2xMS, not forming swollen root hairs as potentially might be expected if the expression of *RSL2* and *RSL3* was elevated in this mutant (Fig. S4). More importantly, the expression levels of *RSL2* and *RSL3* on 2xMS were unaffected in both *obp4* mutants (Fig. S5). Collectively, these results indicate that the decline of *RSL2* and *RSL3* expression on 2xMS is independent from GTL1, DF1 and OBP4.

### GTL1 suppresses *RHD6*, *RSL4* and *LRL3*

Based on our gene expression analysis, *RHD6*, *RSL4* and *LRL3* levels were elevated in *gtl1-1 df1-1* on at least one of the three types of MS media (Fig. 5). Thus, we hypothesized that *RSL4*, *RHD6* and *LRL3* are regulated by GTL1. In order to test this hypothesis, we used a promoter-luciferase assay employing MM2d, an Arabidopsis culture cell line. Using this approach, we found that *GTL1* overexpression strongly reduced *RHD6*, *RSL4* and *LRL3* promoter activities compared to the control (empty) vector (Fig. 6a,b), demonstrating that *RHD6*, *RSL4* and *LRL3* are under the control of GTL1.

**Fig. 6.**
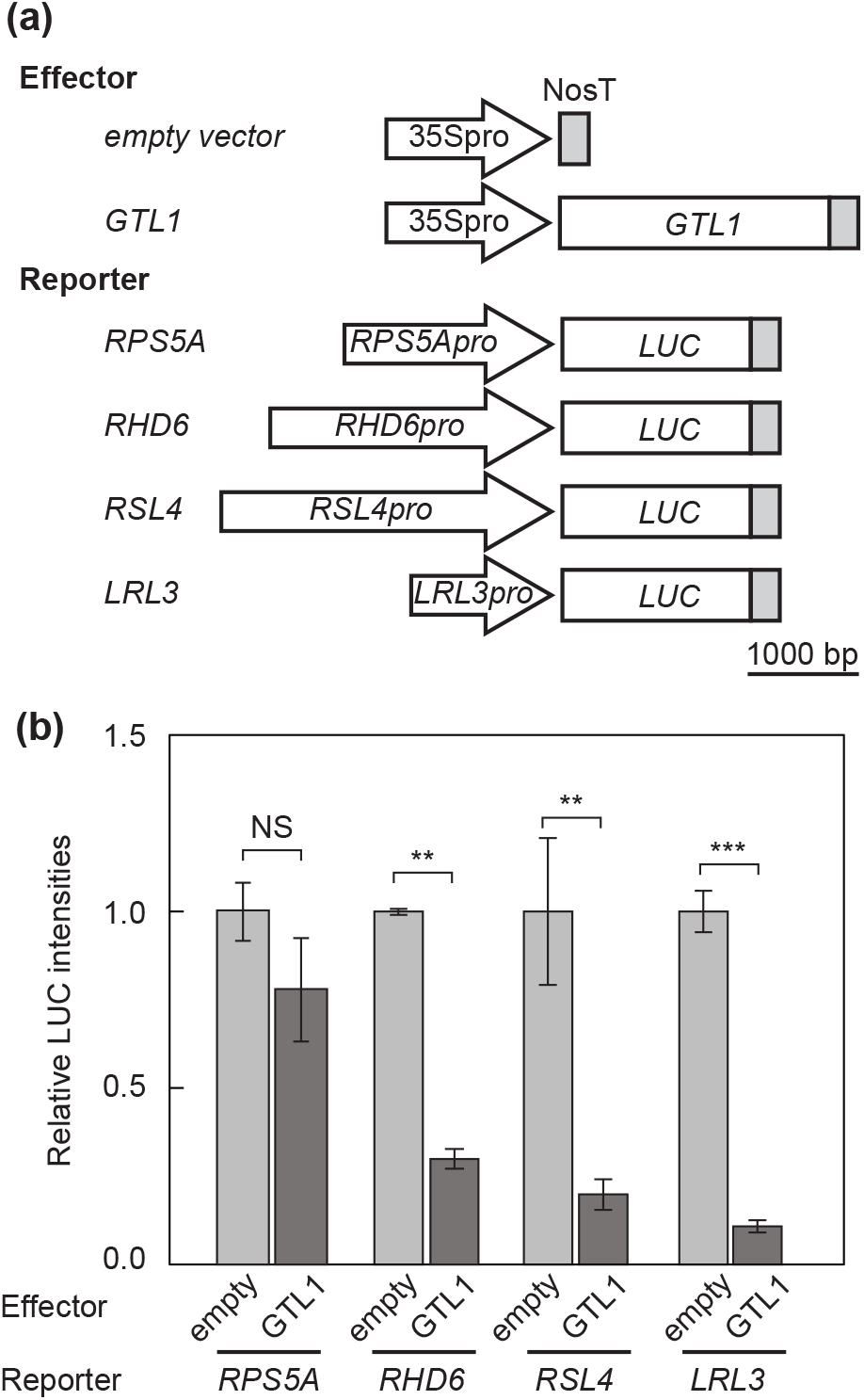
GTL1 suppresses *RHD6*, *RSL4* and *LRL3* expression. (a) Constructs used for the promoter-Luciferase assay. (b) Results from the promoter-Luciferase assay using MM2d culture cells. GTL1 was used as the effector. *RPS5A*, *RHD6*, *RSL1*, or *RSL4* promoter sequences fused with Luciferase were used as reporters. *RPS5A* served as the negative control. Data are mean +/- SD (n = 3). Asterisks indicate significant difference compared to vector control (Student’s t-test, *p < 0.05, **p < 0.01, ***p < 0.001)

Next, since *RSL4* is known as a direct target of RHD6 and GTL1 (Yi *et al*., 2010; Shibata *et al*., 2018), we studied the relationship between GTL1 and RHD6 on *RSL4* promoter activity. As shown in Fig. 7(a), we reproduced that RHD6 and GTL1 work as an activator and a repressor of *RSL4* promoter activity, respectively, in culture cells, consistent with their effects on *RSL4* expression *in planta* (Yi *et al*., 2010; Shibata *et al*., 2018). When both of the transcription factors are introduced together into culture cells, we found that GTL1 dominantly represses *RSL4* promoter activity (Fig. 7a). To identify the regions within the *RSL4* promoter important for GTL1- and RHD6-mediated regulation, a truncated series of *RSL4* promoters were used in another promoter-luciferase assay. When we used a 1500bp region directly upstream from the start codon of *RSL4*, both RHD6 and GTL1 altered *pRSL4*:*LUC* activity (Fig. 7b,c). However, when the fragment was shortened to contain only 500bp immediately upstream of the *RSL4* start codon, RHD6 no longer activated *pRSL4*:*LUC*, while GTL1 still repressed expression of the reporter (Fig. 7b,c). These results indicate that RHD6 and GTL1 bind different regions of the *RSL4* promoter and regulate its expression independently.

**Fig. 7.**
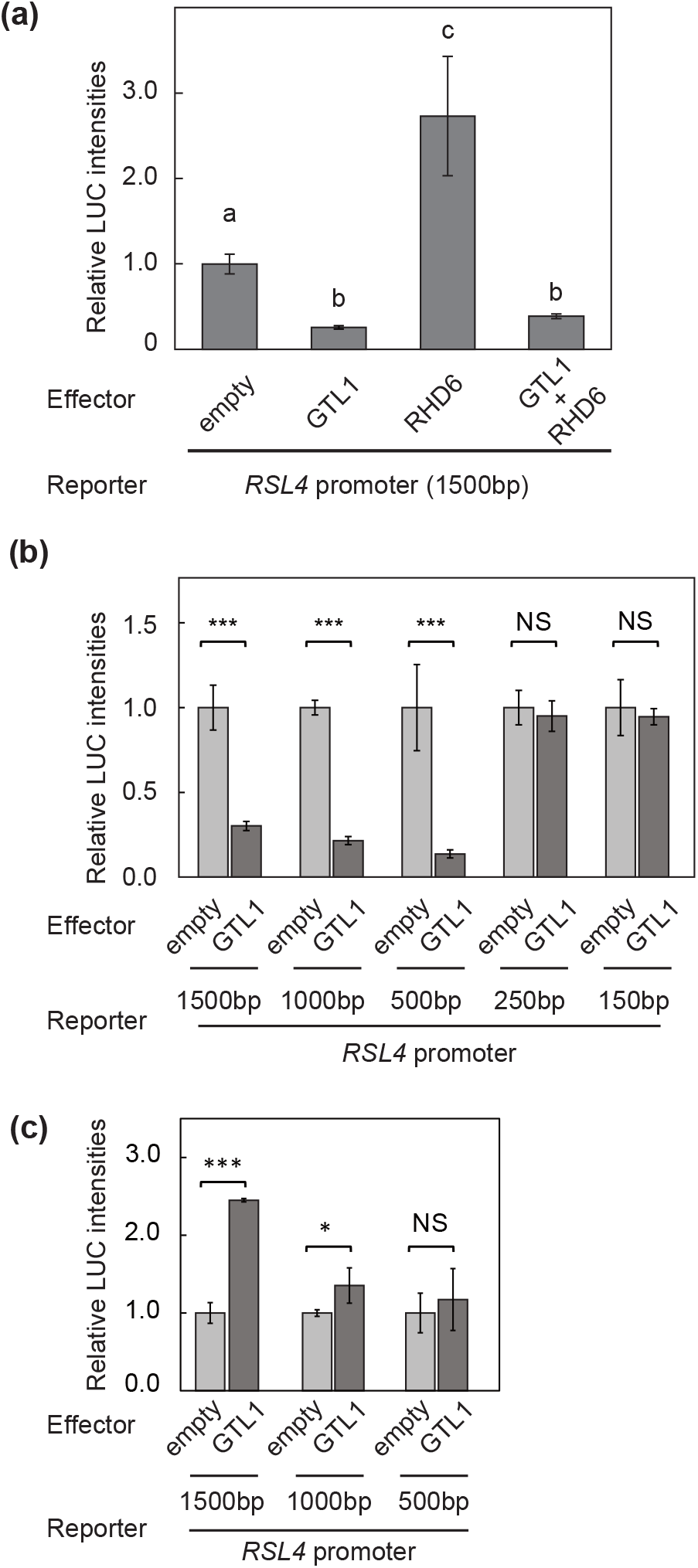
GTL1 dominantly represses *RSL4*. (a) *RHD6* and *GTL1* were both used as effectors in an MM2d-based promoter-Luciferase assay. The *RSL4* promoter region (1500 bp upstream from the start codon) fused with Luciferase was used as a reporter. Different letters indicate significant differences (One-way ANOVA with post-hoc Tukey HSD test, p < 0.01) compared to the control vector (empty). (b,c) Promoter-Luciferase assay using a series of truncated *RSL4* promoter fragments. GTL1 and RHD6 were used as effectors in (b) and (c), respectively. Data are mean +/- SD (n = 3). Asterisks indicate significant differences compared to vector control (Student’s t-test, *p < 0.05, **p < 0.01, ***p < 0.001)

### Ectopic expression of *RSL4* induces abnormal root hairs

Given that *RHD6*, *RSL4*, and *LRL3* promote root hair growth and are repressed by GTL1/DF1 *in planta*, mis-expression of these genes in *gtl1-1 df1-1* may cause the root hair swelling observed in this mutant on 2xMS. In order to test this hypothesis, we generated *rhd6 gtl1 df1*, *rsl4 gtl1 df1* and *lrl3 gtl1 df1* triple mutants. Then, root hair phenotypes were examined on 2xMS.

For *RHD6*, *rhd6-3* is almost hairless even under normal conditions (Masucci & Schiefelbein, 1994; Menand *et al*., 2007), while the *rhd6-3 gtl1-1 df1-1* mutant formed tiny root hairs with balloon-like structures on 2xMS (Fig. 8a), suggesting that RHD6 is not responsible for the phenotype observed in *gtl1-1 df1-1*. In order to further investigate if ectopic expression of *RHD6* causes root hair swelling on 2xMS, we used an *RHD6*-inducible overexpression line (*35S:XVE>>RHD6*), in which 10 µM 17β-estradiol (17ED) treatment increased the level of *RHD6* mRNA by approximately 180-fold compared to mock treatment with DMSO (Fig. S6). Consistent with the WT phenotype, *35S:XVE>>RHD6* treated with DMSO did not form root hairs on 2xMS (Fig. 8b). On the other hand, *35S:XVE>>RHD6* treated with 17ED on 2xMS produced root hairs with normal morphology, in contrast to the expectation that it might cause swollen hairs to form (Fig. 8b). These results demonstrated that the mis-expression of *RHD6* in *gtl1-1 df1-1* does not cause the swollen-root hair phenotype on 2xMS, but instead that *RHD6* promotes the formation of proper root hairs on 2xMS.

**Fig. 8.**
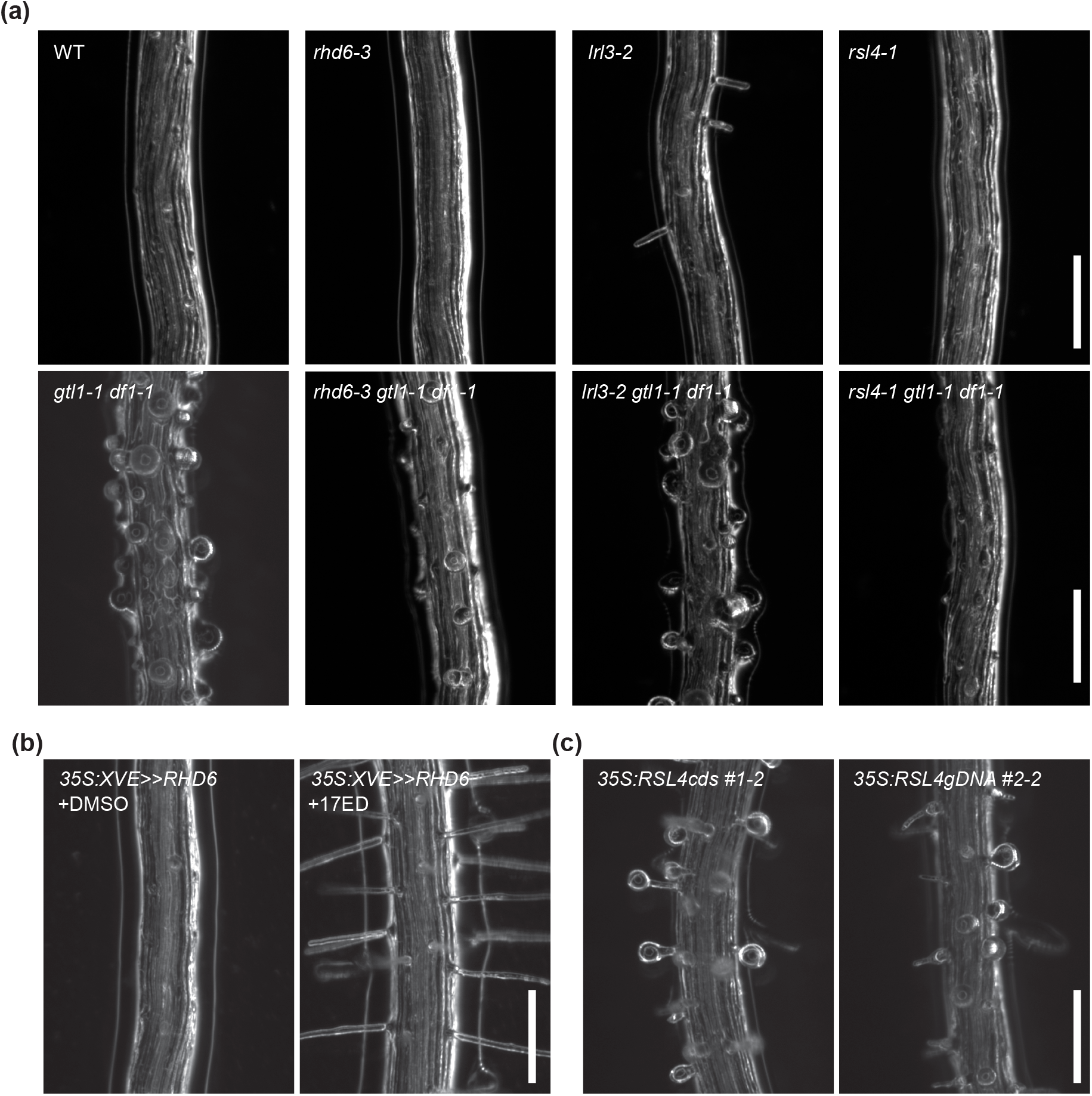
*RSL4* causes aberrant root hair formation on 2xMS. (a) Root hairs of *rhd6-3, lrl3-2, rsl4-1* mutants in the WT (upper panel) or *gtl1-1 df1-1* (lower panel) background grown on 2xMS medium. Note that only the *rsl4* mutation mitigated the root hair swelling phenotype of *gtl1-1 df1-1*. (b) Formation of normal root hairs on 2xMS in plants ectopically over-expressing *RHD6*. DMSO was used as a control treatment. See also Fig. S6. (c) Swollen root hairs in plant ectopically over-expressing *RSL4* on 2xMS. Overexpression of constructs derived from either cDNA (*35S:RSL4cds-GFP*) or the genomic sequence (*35S:gRSL4-GFP*) of RSL4 produced similar phenotypes. Scale bars = 200 µm.

For *LRL3*, the *lrl3-2* single mutant displayed a similar suppression of root hair growth on 2xMS compared to the WT (Fig. 8a, S7). This is consistent with a previous finding that another single mutant for *LRL3*, *lrl3-1*, does not affect root hair growth (Karas *et al*., 2009). Similar to *gtl1-1 df1-1*, *lrl3-2 gtl1-1 df1-1* formed swollen root hairs on 2xMS (Fig. 8a). Therefore, mis-expression of *LRL3* in *gtl1-1 df1-1* does not cause the formation of abnormal root hairs in *gtl1-1 df1-1* on double-strength MS.

In contrast to *RHD6* and *LRL3*, we found that mutation of *RSL4* rescued the *gtl1-1 df1-1* root hair phenotype on 2xMS. (Fig. 8a). Although the *rsl4* single mutant forms root hairs under normal condition due to redundancy with *RSL2* (Shibata *et al*., 2018), both *rsl4-1* and *rsl4-1 gtl1-1 df1-1* exhibited almost no root hair growth on 2xMS. To further investigate if ectopic expression of *RSL4* leads to root hair swelling, we produced lines constitutively overexpressing *RSL4*, using the 35S promoter. These lines exhibited root hair swelling on 2xMS, indicating that mis-expression of *RSL4* causes this phenotype (Fig. 8c). Taken together with the phenotype of the *rsl4-1 gtl1-1 df1-1* triple mutant, these data strongly suggest that mis-expression of *RSL4* in *gtl1-1 df1-1* causes abnormal root hair formation on double-strength MS. Notably, among the *RHD6* subfamily genes, *RSL4* was the only one which we observed to be overexpressed in *gtl1-1 df1-1* on 2xMS (Fig. 5).

## Discussion

Environmental signals affect post-embryonic development of plants. In this study, we showed that excess nutrients, supplied in the form of double-strength MS media, strongly inhibit Arabidopsis growth and have a particularly striking effect on root hairs (Fig. 1). Since one of the main functions of root hairs is to enhance nutrient uptake, the root hair response to 2xMS likely reduces nutrient overloading. Our data further indicate that individual MS components do not account for the effect of 2xMS on root hair growth. Specifically, our custom MS experiments showed that neither the reduction of individual nutrients in 2xMS or their increase in 1/2xMS completely reproduces the root hair phenotypes observed on 1/2xMS or 2xMS, respectively (Fig. 4). These data suggest that Arabidopsis plants integrates signals transduced in response to multiple nutrients and determines an appropriate root hair response.

### GTL1 and DF1 are required for proper root hair response to excess nutrients

We demonstrated that GTL1 and DF1 are critical for suppression of root hair growth on 2xMS, with the *gtl1-1 df1-1* mutant forming frail root hairs on 2xMS (Fig. 2-3). Our AFM analysis showed that both WT and *gtl1-1 df1-1* reduce stiffness of root hairs. However, the reduction of root hair stiffness was more pronounced in *gtl1-1 df1-1* than in the WT (Fig. 3). Consistent with this result, root hair growth of *gtl1-1 df1-1* is hypersensitive to reduction/addition of nutrients (Fig. 4). Additionally, in the *gtl1-1 df1-1* mutant, the expression of *RHD6* subfamily genes was not properly regulated, with some genes, like *RSL4*, up-regulated in this mutant on 2xMS, while others, namely *RSL2* and *RSL3*, were down-regulated (Fig. 5). Thus, we conclude that GTL1 and DF1 function to stabilize changes in gene expression induced by variable environments, thus ensuring appropriate root hair responses.

Notably, the expression levels of *GTL1* and *DF1* gradually decrease with increasing MS concentration, indicating that the strong suppression of root hair growth on 2xMS is not due to induction of *GTL1* and *DF1* (Fig. S3). Instead, nutrient signaling might affect the activities of GTL1 and DF1 via post-translational modifications. Some publicly available proteomic data sets indicate that GTL1 undergoes post-transactional modifications, including phosphorylation and SUMOylation (Reiland *et al*., 2009, 2011; Umezawa *et al*., 2013; Wang *et al*., 2013; Roitinger *et al*., 2015; Choudhary *et al*., 2015). In another study, it was reported that a MAP kinase, MPK4, binds GTL1, although phosphorylation of GTL1 by MPK4 was not detected (Völz *et al*., 2018). In addition, GTL1 activity is known to be regulated by Ca^2+^-dependent calmodulin (Ca^2+^/CaM). Ca^2+^/CaM binds the DNA binding domain of GTL1 and inhibits DNA binding (Weng *et al*., 2012; Yoo *et al*., 2019). Interestingly, it has also been suggested that GTL1 transcriptional-repressor activity is inhibited by Ca^2+^/CaM in response to water-deficit stress, thus leading to derepression of genes related to stomata development (Yoo *et al*., 2010, 2019; Weng *et al*., 2012). These data, together with the lack of transcriptional activation of *GTL1* and *DF1* on 2xMS compared to 1/2xMS, suggest the activity of GTL1 and DF1 is likely regulated by post-translational modification(s) in the context of excess nutrients.

### A GRN enables precise regulation of genes implicated in root hair growth

We demonstrated that WT Arabidopsis plants strongly suppress root hair growth on 2xMS by repressing *RHD6* subfamily genes. Among these genes, at least *RHD6*, *RSL4* and *LRL3* are under the control of GTL1, while the suppression of *RSL2* and *RSL3* appears to be independent of GTL1 and DF1. However, even *RHD6*, *RSL4* and *LRL3* were still responsive to changes in MS concentration in *gtl1-1 df1-1*, suggesting that other transcription factors besides GTL1 and DF1 contribute to repression of these genes (Fig. 5). A C2H2 transcription factor, ZINC-FINGER-PROTEIN1 (ZP1), was recently shown to suppress root hair growth by directly repressing *RHD6*, *RSL2* and *RSL4* (Han *et al*., 2020a). Although this study did not investigate if ZP1 works in the context of environmental responses, it is possible that ZP1 contributes to the repression of root hair growth on 2xMS together with GTL1 and DF1. MYB DOMAIN PROTEIN 30 (MYB30) was also shown recently to negatively regulate root hair growth via direct repression of *RSL4* (Xiao *et al*., 2021). Notably, MYB30 inhibits ETHYLENE INSENSITIVE 3 (EIN3), a positive regulator of ethylene signaling, by directly interacting with this protein (Xiao *et al*., 2021). EIN3, on the other hand, is known to bind RHD6 and promotes root hair growth in response to ethylene signaling (Song *et al*., 2016b). Thus, MYB30 is expected to regulate *RSL4* competitively with EIN3. Additionally, JASMONATE-ZIM-DOMAIN PROTEIN (JAZ) proteins, which are key regulators of jasmonic acid (JA) signaling, also physically interact with RHD6, thus affecting root hair development in response to JA (Han *et al*., 2020b). These data suggest that RHD6 and its binding partners work as a hub within the GRN that regulates root hair development in response to environmental signals. Moreover, our data showed that GTL1 works oppositely to RHD6 in the regulation of *RSL4*. RSL4 itself is also known as a master regulator of root hair growth, promoting growth by directly activating root hair-specific genes (Hwang *et al*., 2017). Interestingly, RSL4 also forms a feedback loop with GTL1 (Shibata *et al*., 2018). Induction of RSL4 activity in glucocorticoid receptor:RSL4 (GR:RSL4)-expressing plants showed that RSL4 induces *GTL1* expression (Vijayakumar *et al*., 2016). On the other hand, we previously demonstrated that GTL1 directly suppresses *RSL4* expression (Shibata *et al*., 2018), similar to the negative effect of GTL1 on *RSL4* observed in this study (Figs. 5-7). Evaluation of the biological significance of this feed-back-loop between GTL1 and RHD6 requires further analysis, using approaches such as mathematical modeling.

Regarding the relationship between *RHD6* and *RSL4*, we showed that the ectopic overexpression of *RHD6* causes plants to form normal root hairs on 2xMS, while *RSL4* does not have this effect (Fig. 8b,c). These results indicate that RHD6 and RSL4 promote the formation of robust and frail root hairs, respectively, on 2xMS. As RHD6 works upstream of *RSL4*, our data suggest that coordinated induction of other TFs by RHD6 in addition to *RSL4*, such as *RSL2*, is important for proper root hair growth in this condition. Since in general plant responses to environmental signals are tightly controlled by a GRN consisting of multiple TFs (Song *et al*., 2016a; Van den Broeck *et al*., 2020), the regulatory network revealed in this study likely fine tunes root hair growth in response to fluctuating environmental conditions.

### Accession numbers

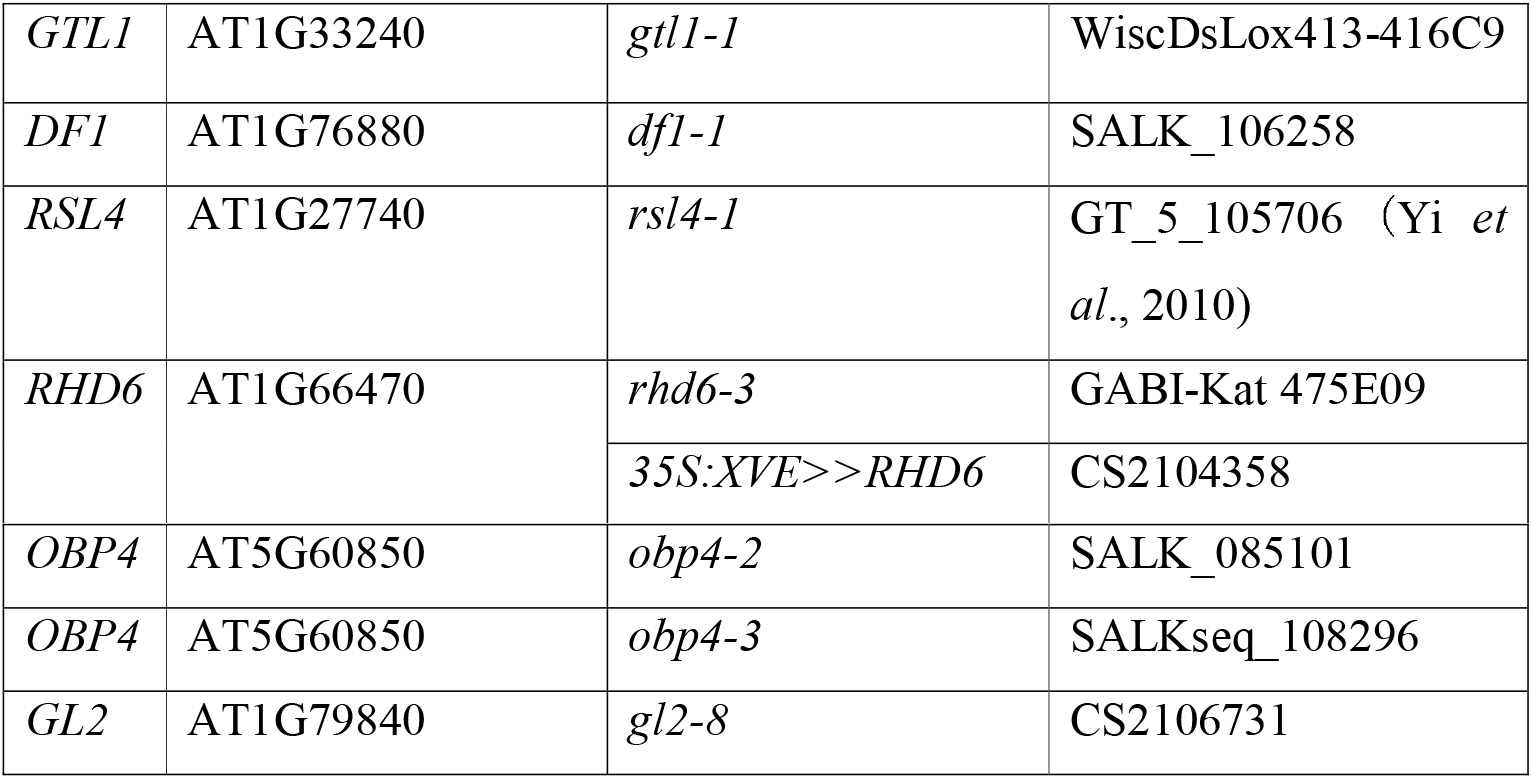

## Acknowledgments

We are grateful to the members of the Sugimoto lab for fruitful discussions about the manuscript. We thank Akiko Hanada, Mariko Mouri, Chika Ikeda and Noriko Doi for their technical assistance. We thank Kazunori Okano and Eri Akita for their technical support with the AFM analysis. We thank Dr. June-Sik Kim for advice regarding statistical tests. This work was supported by grants from MEXT KAKENHI Grant-in-Aid for JSPS Fellows (16J07464) and Grant-in-Aid for Young Scientists (20K15827) to M.S., Grant-in-Aid for JSPS Fellows (19F19781) to D.S.F., Grant-in-Aid for Scientific Research on Priority Areas “Plant-Structure Optimization Strategy” (JP18H05493) to Y.H. and Grant-in-Aid for Transformative Research Areas (A) “Plant Resilience under Fluctuating Environment” (20H05911) to K.S.

## Author contribution

M.S., D.S.F., B.R. and K.S. conceived the research and designed the experiments. M.S., and A.K. performed most of the experiments. R.T., and Y.H. performed AFM analysis. M.S., D.S.F., Y.H. and K.S. wrote the manuscript with input from all co-authors.

## Supplemental Data

**Fig. S1. The expression levels of *EXPA7* and *GL2***

RT-qPCR analysis of *EXPA7* and *GL2* in WT and *gtl1-1 df1-1* grown on 1/2x, 1x and 2x MS media. Expression levels are normalized to that of the *HEL* gene. Data are mean +/- SD. (n = 3, biological replicates). Asterisks indicate a significant difference for the same genotype grown on different types of media (Student’s t-test, *p < 0.05, **p < 0.01, ***p < 0.001).

**Fig. S2. Measurement of root and root hair stiffness**

(a) Force-indentation curves in the AFM measurement.

(b) Box plots showing main root stiffness scores, as determined from analysis of the force indentation curves. Y axis indicates Young’s modulus (*E*). Asterisks indicate a significant difference (Student’s t-test, *p < 0.05, **p < 0.01, ***p < 0.001).

**Fig. S3. The expression levels of *GTL1* and *DF1***

RT-qPCR analysis of *GTL1* and *DF1* grown on 1/2x, 1x and 2x MS media. Expression levels are normalized to that of the *HEL* gene. Data are mean +/- SD. (n = 3, biological replicates). Different letters indicate significant differences for the same genotype grown on different types of media (One-way ANOVA with post-hoc Tukey HSD test, p < 0.05).

**Fig. S4. Roots of *obp4* mutants grown on MS dilution series**

Images of root tips of the WT, *obp4-2* and *obp4-3*, on half- (1/2x), full- (1x) or double- (2x) strength MS medium. Scale bars = 1000 µm

**Fig. S5. The expression levels of *RSL2* and *RSL3* in *obp4* mutants**

RT-qPCR analysis of *RSL2* and *RSL3* grown on 1/2x, 1x and 2x MS media. Expression levels are normalized to that of the *HEL* gene. Data are mean +/- SD. (n = 3, biological replicates). Asterisks indicate a significant difference compared to the WT grown on the same type of medium (Student’s t-test, *p < 0.05, **p < 0.01, ***p < 0.001).

**Fig. S6. The expression levels of *RHD6* and *RSL4* in corresponding overexpression lines**

(a) RT-qPCR analysis of *RHD6* in *35S:XVE>>RHD6*. Total RNA was purified after 24h of treatment with 10μM 17ED. DMSO was used as a control treatment. Expression levels are normalized to that of the *UBQ10* gene. Data are mean +/- SD. (n = 3, biological replicates). Asterisks indicate a significant difference compared to the control condition (Student’s t-test, *p < 0.05, **p < 0.01, ***p < 0.001).

(b) RT-qPCR analysis of *RSL4* in *35S:RSL4cds #1-2 and 35S:gRSL4 #2-2.* Expression levels are normalized to that of the *UBQ10* gene. Data are mean +/- SD. (n = 3, biological replicates). Asterisks indicate a significant difference compared to the WT grown on the same type of medium (Student’s t-test, *p < 0.05, **p < 0.01, ***p < 0.001).

**Fig. S7. Properties of the *LRL3* knock-down mutant**

(a) Gene structure of *AT5G58010*/*LRL3*. Closed boxes denote exons. The triangle represents the position of the T-DNA insertion (SALK_012380/*lrl3-2*). Arrows indicate primers used for genotyping.

(b) Image of an agarose gel showing PCR-amplified DNA fragments for genotyping. Primer sets are shown above each lane. L indicates 100bp Ladder.

(c) Relative expression level of *LRL3* in the *lrl3-2* mutant.

(d) Nucleotide sequence of the genomic region containing the *LRL3* gene. The region between start codons of *LRL3* and *AT5G58020* was used as the *LRL3* promoter in this study.

The sequence and splicing data were obtained from the TAIR database (http://www.arabidopsis.org).

**Table S1. Components of 1/2x, 1x and 2x MS media**

**Table S2. Components of the custom MS media**

**Table S3. List of PCR Primers used in this study**

