## Supplemental Figures for "GTL1 is required for a robust root hair growth response to avoid nutrient overloading"

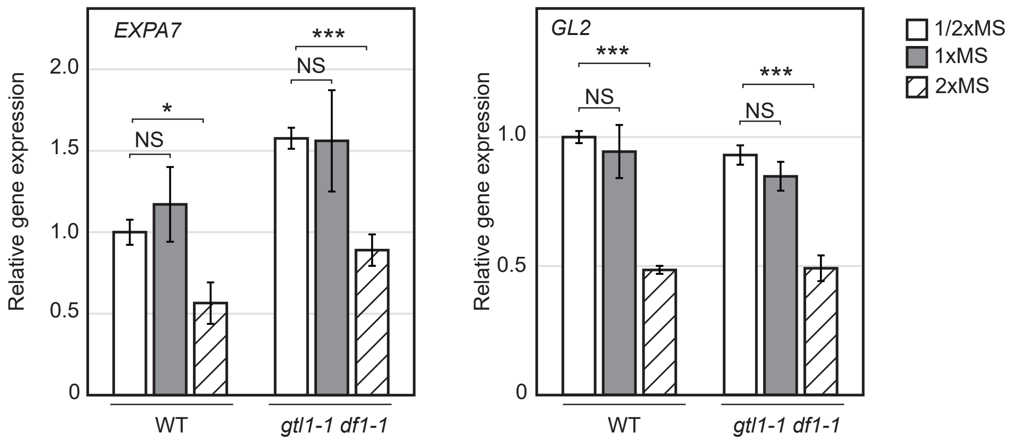


Fig. S1. The expression levels of *EXPA7* and *GL2*

RT-qPCR analysis of *EXPA7* and *GL2* in WT and *gtl1-1 df1-1* grown on 1/2x, 1x and 2x MS media. Expression levels are normalized to that of the *HEL* gene. Data are mean +/- SD. (n = 3, biological replicates). Asterisks indicate a significant difference for the same genotype grown on different types of media (Student' s t-test, *p < 0.05, **p < 0.01, ***p < 0.001).

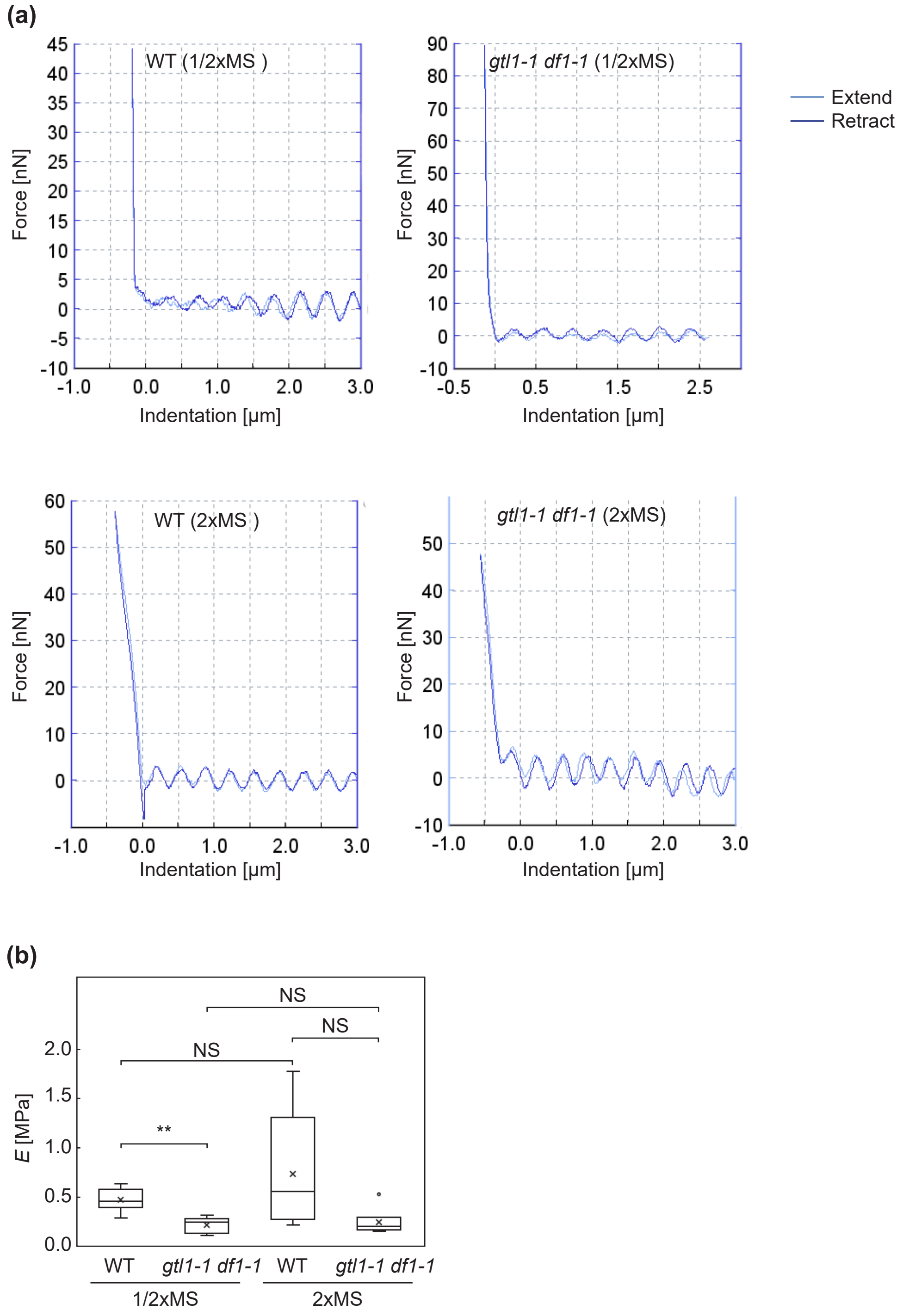


Fig. S2. Measurement of root and root hair stiffness

(a) Force-indentation curves in the AFM measurement.

(b) Box plots showing main root stiffness scores, as determined from analysis of the force indentation curves. Y axis indicates Young's modulus (E). Asterisks indicate a significant difference (Student' s t-test, *p < 0.01, **p < 0.01, ***p < 0.001).

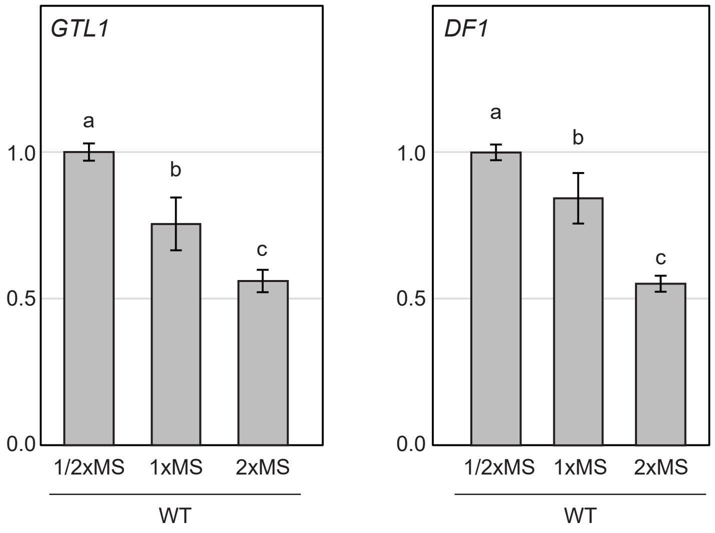


Fig. S3. The expression levels of *GTL1* and *DF1*

RT-qPCR analysis of *GTL1* and *DF1* grown on 1/2x, 1x and 2x MS media. Expression levels are normalized to that of the *HEL* gene. Data are mean +/- SD. (n = 3, biological replicates). Different letters indicate significant differences for the same genotype grown on different types of media (One-way ANOVA with post-hoc Tukey HSD test, $p < 0.05$).

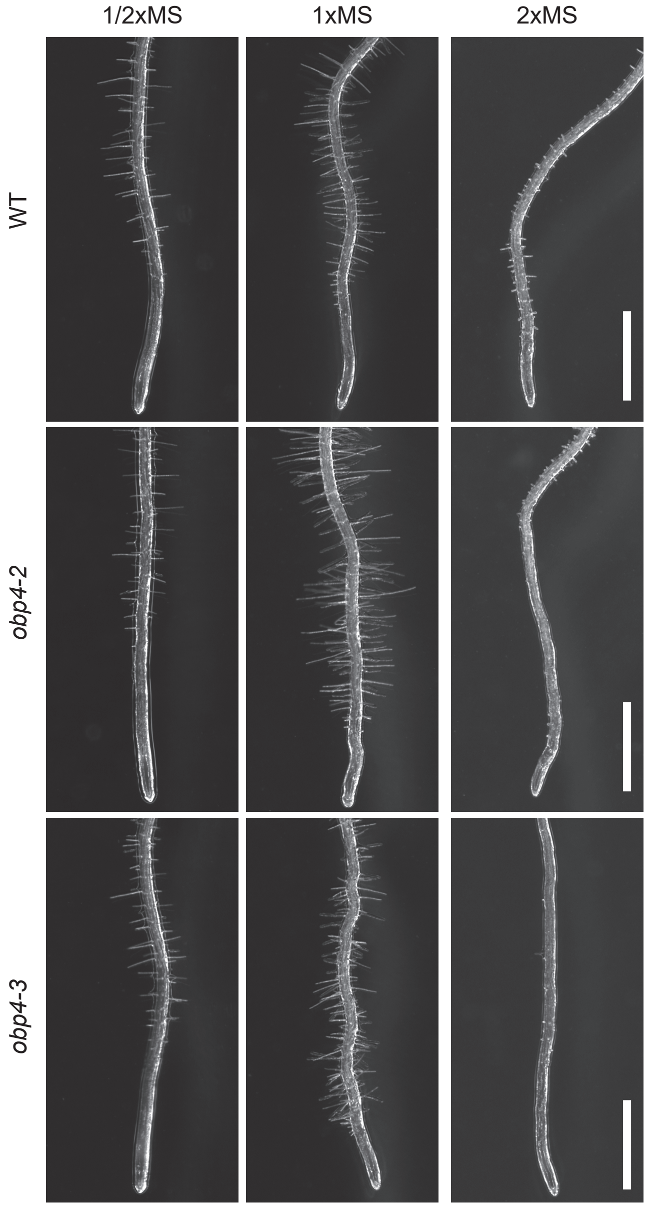


Fig. S4. The roots of *obp4* mutants grown on MS dilution series

Images of root tips of the WT, *obp4-2* and *obp4-3*, on half- (1/2 x), full- (1 x) or double- (2 x) strength MS medium. Scale bars = 1000 μm

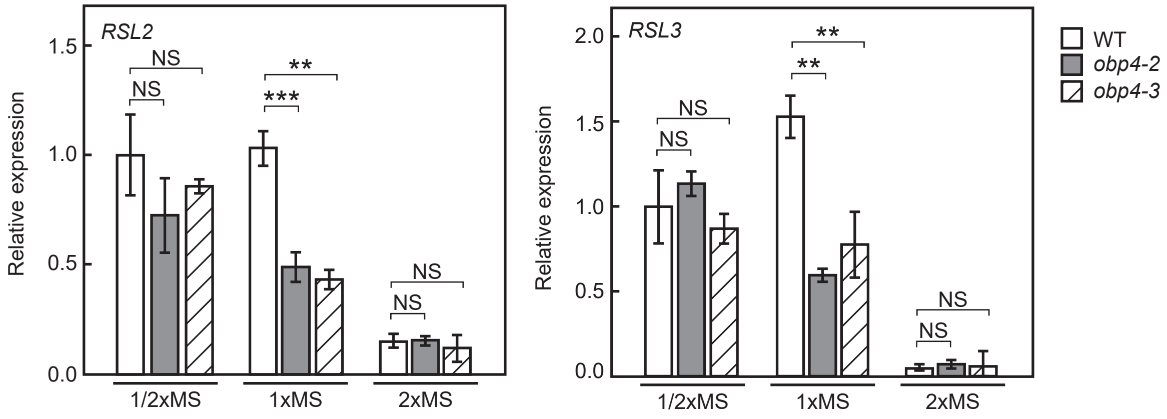


Fig. S5. The expression levels of *RSL2* and *RSL3* in *obp4* mutants

RT-qPCR analysis of *RSL2* and *RSL3* grown on 1/2x, 1x and 2x MS media. Expression levels are normalized to that of the *HEL* gene. Data are mean +/- SD. (n = 3, biological replicates). Asterisks indicate a significant difference compared to the WT grown on the same type of medium (Student' s t-test, *p < 0.05, **p < 0.01, ***p < 0.001).

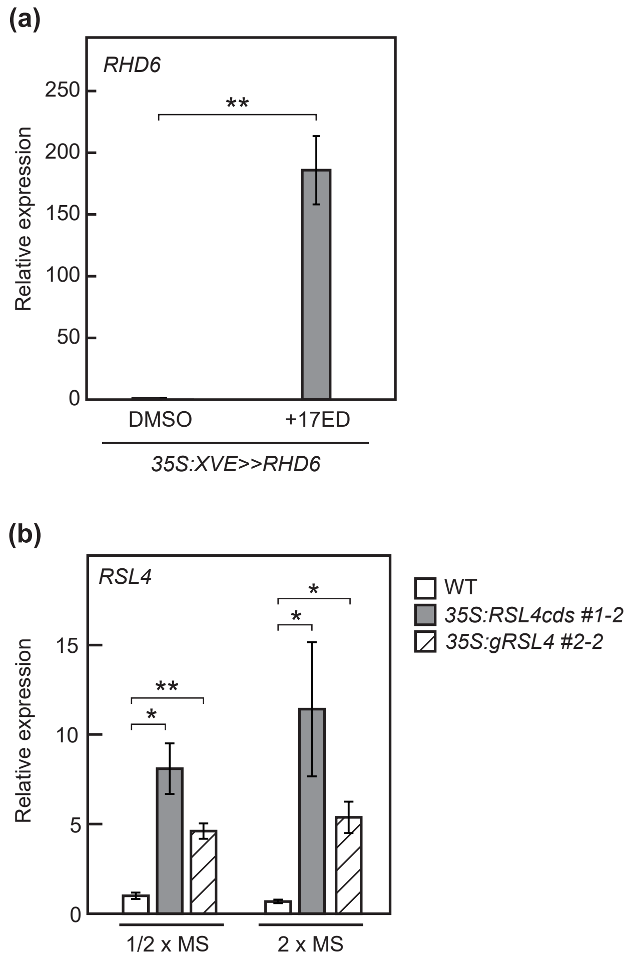


Fig. S6. The expression levels of *RHD6* and *RSL4* in corresponding overexpression lines

(a) RT-qPCR analysis of *RHD6* in *35S:XVE>>RHD6*. Total RNA was purified after 24h of treatment with 10μM 17ED. DMSO was used as a control treatment. Expression levels are normalized to that of the *UBQ10* gene. Data are mean +/- SD. (n = 3, biological replicates). Asterisks indicate a significant difference compared to the control condition (Student' s t-test, *p < 0.05, **p < 0.01, ***p < 0.001).

(b) RT-qPCR analysis of *RSL4* in *35S:RSL4cds #1-2* and *35S:gRSL4 #2-2*. Expression levels are normalized to that of the *UBQ10* gene. Data are mean +/- SD. (n = 3, biological replicates). Asterisks indicate a significant difference compared to the WT grown on the same type of medium (Student' s t-test, *p < 0.05, **p < 0.01, ***p < 0.001).

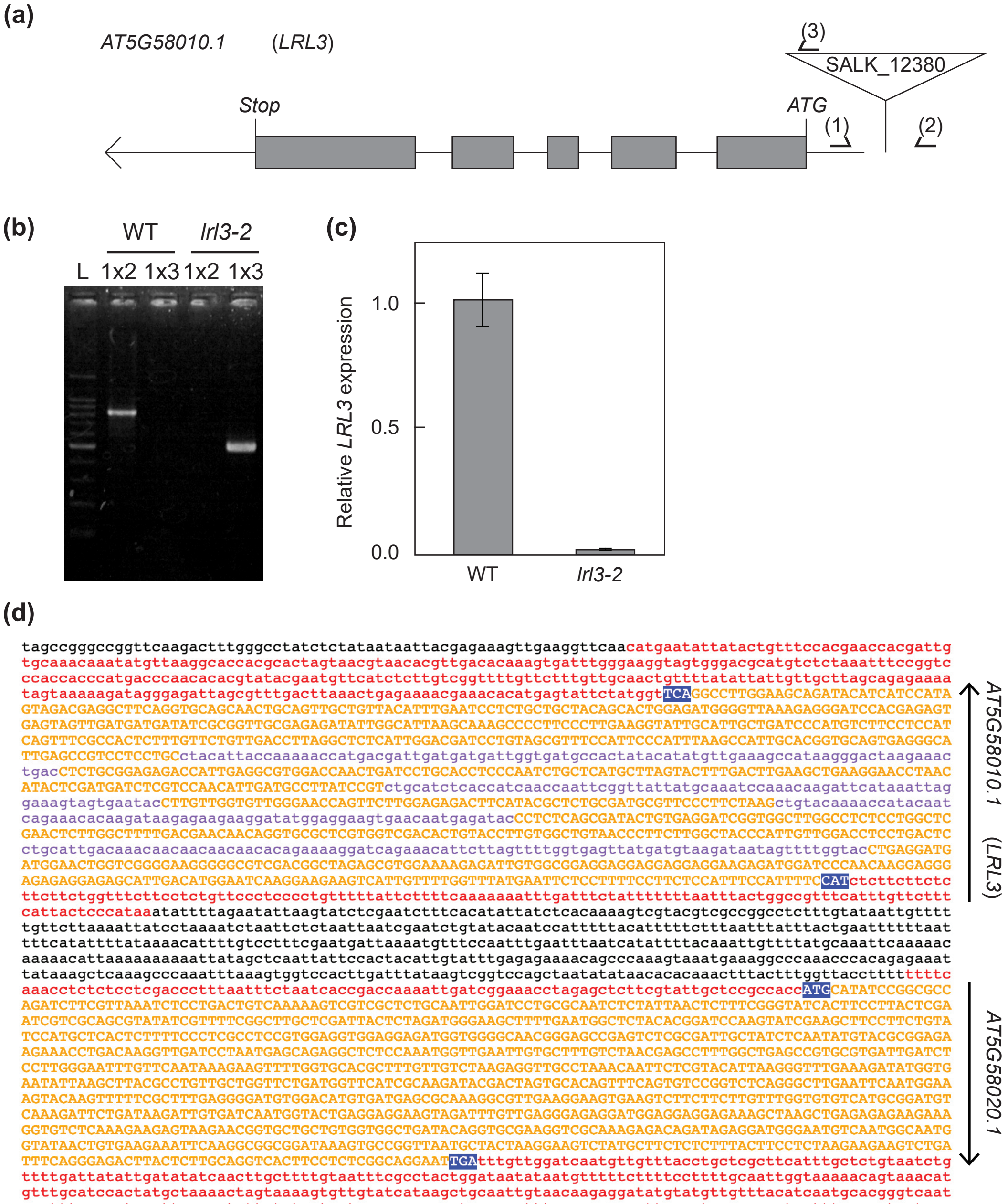


Fig. S7. Properties of the *LRL3* knock-down mutant

(a) Gene structure of *AT5G58010*/*LRL3*. Closed boxes denote exons. The triangle represents the positions of the T-DNA insertion (SALK_012380/*lrl3-2*). Arrows indicate primers used for genotyping.

(b) Image of an agarose gel showing PCR-amplified DNA fragments for genotyping. Primer sets are shown above each lane. L indicates 100bp Ladder.

(c) Relative expression level of *LRL3* in the *lrl3-2* mutant.

(d) Nucleotide sequence of the genomic region containing *LRL3* gene. The region between start codons of *LRL3* and *AT5G58020* was used as the *LRL3* promoter in this study.

The sequence and splicing data were obtained from the TAIR database (http://www.arabidopsis.org).
