## Supplementary material for "GTL1 is required for a robust root hair growth response to avoid nutrient overloading": Table S1

Table S1. Components of 1/2x, 1x and 2x MS media

| chemicals | 1/2xMS [mg/l] | 1xMS [mg/l] | 2xMS [mg/l] |
|---|---|---|---|
| $KNO_3$ | 950 | 1900 | 3800 |
| $NH_4NO_3$ | 825 | 1650 | 3300 |
| $CaCl_2 \cdot 2H_2O$ | 220 | 440 | 880 |
| MgSO4·7H2O | 185 | 370 | 740 |
| $KH_2PO_4$ | 85 | 170 | 340 |
| 2NaEDTA 2H2O | 18.65 | 37.3 | 74.6 |
| $FeSO_4 \cdot 7H_2O$ | 13.9 | 27.8 | 55.6 |
| MnSO4·5H2O | 12.05 | 24.1 | 48.2 |
| $ZnSO_4 \cdot 7H_2O$ | 4.3 | 8.6 | 17.2 |
| $H_3BO_3$ | 3.1 | 6.2 | 12.4 |
| KI | 0.415 | 0.83 | 1.66 |
| $Na_2MoO_4 \cdot 2H_2O$ | 0.125 | 0.25 | 0.5 |
| CuSO4·5H2O | 0.0125 | 0.025 | 0.05 |
| $CoCl_2 \cdot 6H_2O$ | 0.0125 | 0.025 | 0.05 |

| | |
|---|---|
| sucrose (g) | 10 |
| 250mM MES pH5.7 (ml) | 10 |
| 55mM myo-inositol (µl) | 1000 |
| gellan gum (g) | 6 |
