## Supplementary material for "GTL1 is required for a robust root hair growth response to avoid nutrient overloading": Table S2

Table S2. Components of the custom MS media

| omission of nutrient composition from 2xMS media | | | | | | | | | | | | | | |
|---|---|---|---|---|---|---|---|---|---|---|---|---|---|---|
| | $KNO_3$ | $NH_4NO_3$ | $CaCl_2·2H_2O$ | MgSO4·7H2O | $KH_2PO_4$ | 2NaEDTA 2H2O | $FeSO_4·7H_2O$ | MnSO4·5H2O | $ZnSO_4·7H_2O$ | $H_3BO_3$ | KI | $Na_2MoO_4·2H_2O$ | CuSO4·5H2O | $CoCl_2·6H_2O$ |
| 2xMS_All | 2x | 2x | 2x | 2x | 2x | 2x | 2x | 2x | 2x | 2x | 2x | 2x | 2x | 2x |
| 2xMS_1/2x$KNO_3$ | 1/2x | 2x | 2x | 2x | 2x | 2x | 2x | 2x | 2x | 2x | 2x | 2x | 2x | 2x |
| 2xMS_1/2x$NH_4NO_3$ | 2x | 1/2x | 2x | 2x | 2x | 2x | 2x | 2x | 2x | 2x | 2x | 2x | 2x | 2x |
| 2xMS_1/2x$CaCl_2$ | 2x | 2x | 1/2x | 2x | 2x | 2x | 2x | 2x | 2x | 2x | 2x | 2x | 2x | 2x |
| 2xMS_1/2x$MgSO_4$ | 2x | 2x | 2x | 1/2x | 2x | 2x | 2x | 2x | 2x | 2x | 2x | 2x | 2x | 2x |
| 2xMS_1/2x$KH_2PO_4$ | 2x | 2x | 2x | 2x | 1/2x | 2x | 2x | 2x | 2x | 2x | 2x | 2x | 2x | 2x |
| 2xMS_1/2x$FeSO_4$ 1/2xEDTA | 2x | 2x | 2x | 2x | 2x | 1/2x | 1/2x | 2x | 2x | 2x | 2x | 2x | 2x | 2x |
| 2xMS_1/2x$MnSO_4$ | 2x | 2x | 2x | 2x | 2x | 2x | 2x | 1/2x | 2x | 2x | 2x | 2x | 2x | 2x |
| 2xMS_1/2x$ZnSO_4$ | 2x | 2x | 2x | 2x | 2x | 2x | 2x | 2x | 1/2x | 2x | 2x | 2x | 2x | 2x |
| 2xMS_1/2x$H_3BO_3$ | 2x | 2x | 2x | 2x | 2x | 2x | 2x | 2x | 2x | 1/2x | 2x | 2x | 2x | 2x |
| 2xMS_1/2xKI | 2x | 2x | 2x | 2x | 2x | 2x | 2x | 2x | 2x | 2x | 1/2x | 2x | 2x | 2x |
| 2xMS_1/2x$Na_2MoO_4$ | 2x | 2x | 2x | 2x | 2x | 2x | 2x | 2x | 2x | 2x | 2x | 1/2x | 2x | 2x |
| 2xMS_1/2xCuSO | 2x | 2x | 2x | 2x | 2x | 2x | 2x | 2x | 2x | 2x | 2x | 2x | 1/2x | 2x |
| 2xMS_1/2x$CoCl_2$ | 2x | 2x | 2x | 2x | 2x | 2x | 2x | 2x | 2x | 2x | 2x | 2x | 2x | 1/2x |

| addition of nutrient composition from 1/2xMS media | | | | | | | | | | | | | | |
|---|---|---|---|---|---|---|---|---|---|---|---|---|---|---|
| | $KNO_3$ | $NH_4NO_3$ | $CaCl_2·2H_2O$ | MgSO4·7H2O | $KH_2PO_4$ | 2NaEDTA 2H2O | $FeSO_4·7H_2O$ | MnSO4·5H2O | $ZnSO_4·7H_2O$ | $H_3BO_3$ | KI | $Na_2MoO_4·2H_2O$ | CuSO4·5H2O | $CoCl_2·6H_2O$ |
| 1/2xMS_All | 1/2x | 1/2x | 1/2x | 1/2x | 1/2x | 1/2x | 1/2x | 1/2x | 1/2x | 1/2x | 1/2x | 1/2x | 1/2x | 1/2x |
| 1/2xMS_2x$KNO_3$ | 2x | 1/2x | 1/2x | 1/2x | 1/2x | 1/2x | 1/2x | 1/2x | 1/2x | 1/2x | 1/2x | 1/2x | 1/2x | 1/2x |
| 1/2xMS_2x$NH_4NO_3$ | 1/2x | 2x | 1/2x | 1/2x | 1/2x | 1/2x | 1/2x | 1/2x | 1/2x | 1/2x | 1/2x | 1/2x | 1/2x | 1/2x |
| 1/2xMS_2x$KNO_3$ 2xNH4$NO_3$ | 2x | 2x | 1/2x | 1/2x | 1/2x | 1/2x | 1/2x | 1/2x | 1/2x | 1/2x | 1/2x | 1/2x | 1/2x | 1/2x |
| 1/2xMS_2x$FeSO_4$+2xEDTA | 1/2x | 1/2x | 1/2x | 1/2x | 1/2x | 2x | 2x | 1/2x | 1/2x | 1/2x | 1/2x | 1/2x | 1/2x | 1/2x |
