## Supplementary material for "GTL1 is required for a robust root hair growth response to avoid nutrient overloading": Table S3-1

Table S3. List of PCR Primers used in this study

Promoter cloning for LUC assay

| | Primer name | Sequence |
|---|---|---|
| *RSL4 promoter* | SacII-RSL4pro-1500-F | ATACCGCGGAGGTGCAATGTACCGTAACC |
| | BamHI-RSL4pro-R | ATAGGATCCCGCTCTAACTGATCAACTCTTG |
| | HindIII-RSL4p-1000-F | TCAGTAAGCTTGTGTTGAATTCGCCTTATTTGC |
| | HindIII-RSL4p-500-F | TCAGTAAGCTTTGATTCCACAAATCGTGCAT |
| | HindIII-RSL4p-250-F | TCAGTAAGCTTCATGCATGGCTTCGTTTCAC |
| | HindIII-RSL4p-150-F | TCAGTAAGCTTCATCACCAAATCTTCCTTGAG |
| | HindIII-pGEM-R | ATCATAAGCTTCCTATAGTGAGTCGTATTAC |
| *LRL3 promoter* | SacII-LRL3-pro603-F | ATACCGCGGAGGTAACCAAAGTAAAGTTTGTGTGTT |
| | BamHI-LRL3-proR | ATAGGATCCCTCTTCTTCTCTTCTTCTGGTTTC |
| *RHD6 promoter* | RHD6 proF | AAAGAATGGGCCGAATGTC |
| | RHD6 proR | TAGACACTAATAAGTTTGATAAGTGA |
| *RSL1 promoter* | SacII RSL1pro Fw | AAAGTAGCTAACCCGCGGTGGACATTTGCTTTACTTGTGG |
| | BamHI RSL1pro RV | AAATTAAAGCTAACGGATCCTGGTACTAAAGGGTGTTCTAGTG |
| *RSL4 promoter* | SacII RSL4pro Fw | AAAGTAGCTAACCCGCGGGTGTGTGCATGCATGTGTGT |
| | BamHI RSL4pro RV | AAATTAAAGCTAACGGATCCCGCTCTAACTGATCAACTCTTGCC |
| *RSL3 promoter* | SacII RSL2L1pro Fw | AAAGTAGCTAACCCGCGGGCTTGAAATATGGCTGCACA |
| | BamHI RLS2L1pro RV | AAATTAAAGCTAACGGATCCTTTTGATCACTAAGCGACTTTAAC |
| *RPS5A promoter* | RSP5Apro-F | GCTTGATAAACTATCGCCACC |
| | RSP5Apro-R | GGCTGTGGTGAGAGAAACAG |

RSL4 cloning for overexpression lines

| | | |
|---|---|---|
| genomic *RSL4* and *RSL4* cDNA | attB1-RSL4-F | GGGGACAAGTTTGTACAAAAAAGCAGGCTGCATGGACGTTTTTGTTGATGGTG |
| | attB2-RSL4_noStop-R | GGGGACCACTTTGTACAAGAAAGCTGGGTCCATAAGCCGAGACAAAAGG |
| | attB2-RSL4_withStop-R | GGGGACCACTTTGTACAAGAAAGCTGGGTCTCACATAAGCCGAGACAAAAGG |
