## Supplementary material for "GTL1 is required for a robust root hair growth response to avoid nutrient overloading": Table S3-2

| mutant name | Accettion | Primer name | Sequence | For T-DNA | Reference |
|---|---|---|---|---|---|
| *gtl1-1* | WiscDsLox413-416C9 | *WF627-F1* | TTCTCGTCTCATAGCTCATCG | 627-R1 x Wis_pDS-LOX-LB | Breuer et al 2009 |
| | | *WF627-R1* | TGGCCATCTTGATGATGATGG | | Breuer et al 2009 |
| | | Wis_pDS-LOX-LB | GGGTTTCGCTCATGTGTTGA | | This study |
| *df1-1* | SALK_106258 | DF1_258-F1 | TGCTGATTGATCCACTTCTCA | 258R5 x LBa1 | Shibata et al 2018 |
| | | DF1_258 R5 | GAGATGTTTGGAAACGAAGGTAC | | Shibata et al 2018 |
| | | LBa1 | GCGTGGACCGCTTGCTGCAACT | | - |
| *obp4-2* | SALKseq_085101 | obp4-2 LP | TCAAGCAACGTAGTTCAATGG | LBb1.3 x RP | Rymen et al., 2017 |
| | | obp4-2 RP | GTTCTTGCAAAAGTGACGAGG | | Rymen et al., 2017 |
| | | LBb1.3 | ATTTTGCCGATTTCGGAAC | | - |
| *obp4-3* | SALKseq_108296 | obp4-3 LP | CCGTTTTAAACCCATCAAATAAAA | LBb1.3 x RP | Rymen et al., 2017 |
| | | obp4-3 RP | TTTACGGTAACGGGATCGAG | | Rymen et al., 2017 |
| | | LBb1.3 | ATTTTGCCGATTTCGGAAC | | - |
| *rhd6-3* | GABI-Kat 475E09 | rhd6 F | GTTCCCAATGGCACCAAGGTACA | rhd6 F x GABI LB | Menand et al 2007 |
| | | rhd6 R | TAACTTGAGAAATGTCAGGAGC | | This study |
| | | GABI LB | CCCATTTGGACGTGAATGTAGACAC | | Menand et al 2007 |
| *rsl4-1* | GT_5_105706 | rsl4-1 homozygote F | GAAAGCTTCGGTCACAAGTGTTAAA | JIC-RB1 x rsl4-1 LB R | Yi et al 2010 |
| | | rsl4-1 LB R | TTGTAAGCCAATGGTGCGTACAT | | Yi et al 2010 |
| | | JIC-RB1 | CCGAACAAAAATACCGGTTCCC | | Yi et al 2010 |
| *Atlrl3(380)* | SALK_012380 | lrl3_380_LP2 | CGAGCAAGCCGAAAACGATA | LBa1 x RP2 | This study |
| | | lrl3_380_RP2 | CTCTGTTCCCTCCCCTGTTT | | This study |
| | | LBa1 | GCGTGGACCGCTTGCTGCAACT | | - |
