## Supplementary material for "GTL1 is required for a robust root hair growth response to avoid nutrient overloading": Table S3-3

| | Primer name | Sequence | Reference |
|---|---|---|---|
| For qPCR of *GTL1* | GTL1-627F2 | ATGGAATTGTTTGAAGGTTTGG | Breuer et al 2009 |
| | GTL1-627R2 | GACATGACCTCGTGTTCTCG | Breuer et al 2009 |
| For qPCR of *DF1* | DF1F5KN | GACATGGGAATAGCGTTTCG | Shibata et al 2018 |
| | DF1R4KN | TCGGCATAACTGTCGTTACC | Shibata et al 2018 |
| For qPCR of *RSL1* | RSL1_FWD | TCGTACCGCTACTCGGCTTCTT | Rymen et al 2017 |
| | RSL1_REV | CAATAAACGGCCTTTCACGGGAGA | Rymen et al 2017 |
| For qPCR of *RSL2* | RSL2_FWD | CTCGTCCCCAATGGAACAAAGGTC | Rymen et al 2017 |
| | RSL2_REV | GCAATCGGCGCATACATCCATAGA | Rymen et al 2017 |
| For qPCR of *RSL3* | RSL3_FWD | TCGTCCCTAATGGAACAAAGGTTG | Rymen et al 2017 |
| | RSL3_REV | GGCCAATGTCCATTCCGTTGTAAG | Rymen et al 2017 |
| For qPCR of *RSL4* | qRSL4-F | AGGCAAAACTAGAGCCACCA | Shibata et al 2018 |
| | qRSL4-R | ATCGACTTTTGTCCCGTTTG | Shibata et al 2018 |
| For qPCR of *RHD6* | qRHD6-F | TCACGAGAGCTTTCCTCCTC | Shibata et al 2018 |
| | qRHD6-R | TGAAGCCGTAGCTCATGTTG | Shibata et al 2018 |
| For qPCR of *LRL1* | qLRL1-F | TCTCAAATCTCCGAGGCTGG | This study |
| | qLRL1-R | TTTGGCCACTTGATGTTCCG | This study |
| For qPCR of *LRL2* | qLRL2-F | GATACCGGCGTTCCTTTGTC | This study |
| | qLRL2-R | AACGGAGGGAGCGTCTTTAA | This study |
| For qPCR of *LRL3* | qLRL3-F | AGATTGGGAGGTGCAGGATC | This study |
| | qLRL3-R | TCCATCAGTTTCGCCACTCT | This study |
| For qPCR of *EXPA7* | qEXPA7-F | TTAACAGCGGCTACGGACTG | This study |
| | qEXPA7-R | GGCAAAGATTGGTGGCTGTG | This study |
| For qPCR of *GL2* | qGL2-F | CCGATGATCTCCACCTCGAA | This study |
| | qGL2-R | GTGTTCTTGATCGTCGGAGC | This study |
| For qPCR of *UBQ10* | UBQ10-F | GAAGTGGAAAGCTCCGACAC | Shibata et al 2018 |
| | UBQ10-R | TTAGAAACCACCACGAAGACG | Shibata et al 2018 |
| For qPCR of *HEL* | HEL_hk_Fw | CCATTCTACTTTTTGGCGGCT | Rymen et al 2017 |
| | HEL_hk_Rv | TCAATGGTAACTGATCCACTCTGATG | Rymen et al 2017 |
